# Chemical control of CSA geometry enables relaxation-optimized ^19^F-^13^C NMR probes

**DOI:** 10.64898/2026.01.19.700399

**Authors:** Jin-Gon Shim, Nikol N. Georgieva, Scott A. Robson, Nikola T. Burdzhiev, Ognian I. Petrov, Jinlei Cui, Anupama Acharya, Ilya Kuprov, Vladimir Gelev, Joshua J. Ziarek

## Abstract

Fluorine NMR is a powerful tool for probing biomolecular structure and dynamics, yet the performance of ^19^F probes is fundamentally constrained by rapid transverse relaxation driven by chemical shift anisotropy (CSA). Despite its central role, CSA has largely been treated as an immutable nuclear property rather than a chemically addressable design parameter. Here we demonstrate that the *geometry* of the CSA tensor – specifically its magnitude, symmetry, and orientation relative to the internuclear dipolar interaction – constitutes a decisive and engineerable determinant of relaxation behavior in coupled ^19^F-^13^C spin systems. Guided by electronic-structure calculations and Bloch-Redfield-Wangsness relaxation theory, we establish quantitative design rules that predict when CSA-dipolar interference can be exploited to suppress transverse relaxation. Implementation of these principles in a cysteine-reactive fluoropyrimidine scaffold yields a reporter that supports simultaneous 19F and 13C TROSY optimization, validated by solid-state MAS NMR and protein-based experiments. When incorporated into the 42 kDa maltose binding protein, the probe exhibits exceptionally slow ^13^C transverse relaxation (R_2_ ≈ 2-3 s^−1^) corresponding to linewidths of ∼2 Hz that persist even at apparent molecular weights exceeding 200 kDa. These results recast relaxation optimization as a chemically programmable problem and provide a general framework for the rational design of next-generation NMR probes tailored to large, dynamic, and heterogeneous biomolecular systems.

## INTRODUCTION

Fluorine NMR has become a powerful approach for probing the structure, dynamics, and energetics of biological macromolecules because the ^19^F nucleus combines high sensitivity with complete biological silence and large chemical-shift dispersion^1^. Cysteine-reactive ^19^F probes have played a central role in this development, enabling site-specific reporting in proteins that are otherwise difficult to interrogate by traditional isotopic labeling strategies. These tags have revealed conformational equilibria in G protein-coupled receptors^2–6^, membrane transporters^7^ and enzymes^8^, providing access to dynamical processes spanning microseconds to seconds. Despite these advances, the underlying probe chemistry remains largely empirical, and fundamental constraints in current designs limit the scope of biological problems accessible by ^19^F NMR. As highlighted by Gronenborn, the behavior of ^19^F nuclei in proteins remains difficult to predict at a fundamental level: no general theoretical framework exists for fluorine chemical shifts, and quantitative interpretation of ^19^F relaxation is still constrained by the scarcity of experimentally benchmarked chemical shift anisotropy (CSA) tensors and transferable relaxation parameters^1,9^. Consistent with this view, recent relaxation studies of monofluorinated tryptophans demonstrated that ^19^F transverse relaxation in aromatic systems is dominated by CSA and that CSA-governed behavior observed in free amino acids transfers reliably to their protein-bound counterparts^10^. Together, these findings underscore the need for systematic CSA characterization and predictive design strategies that move beyond empirical probe optimization.

The most widely used cysteine-reactive tags – trifluoromethyl-based probes such as TET and BTFMA – are conceptually limited in several respects^11^. First, they provide only a single ^19^F nucleus, precluding heteronuclear correlations and multidimensional spectroscopic assignments. As a result, analyses rely almost exclusively on one-dimensional line shapes, which must be deconvolved into substate populations using user-defined models that introduce substantial bias and ambiguity^12^. Second, the intrinsically large and untunable chemical-shift anisotropy (CSA) of CF_3_ groups produces rapid transverse relaxation and broad lines, especially in large or slowly tumbling proteins^13^. This relaxation penalty becomes severe at high magnetic field strengths, where linewidths scale strongly with CSA magnitude. Third, rapid threefold rotation of the CF_3_ group leads to motional averaging of local shielding interactions, reducing sensitivity to subtle conformational or environmental differences^14^.

Recent efforts have sought to overcome aspects of these constraints by redesigning the ^19^F chemical environment to enhance either conformational sensitivity or environmental responsiveness. Environmentally ultrasensitive monofluoroethyl (mFE) tags^15^ and electronically tuned fluorinated pyridone probes^16^ have expanded the available chemical-shift range, enabling detection of low-population states and improved discrimination of conformational ensembles through amplified isotropic ¹⁹F chemical-shift responses. Yet, these approaches do not address the core physical limitation that dominates spectral quality in large biomolecules: the inability to rationally engineer transverse relaxation through control of CSA geometry. No existing probe family provides a systematic route to modulate CSA magnitude, symmetry, or orientation to favor destructive CSA-dipolar interference, nor do they incorporate a directly bonded heteronucleus that enables dual-nucleus TROSY or multidimensional correlations^17^.

Here we address these limitations by establishing a generalizable design framework for relaxation-optimized ^19^F-^13^C spin systems, guided by electronic-structure theory and Bloch–Redfield–Wangsness (BRW) relaxation analysis. We show that substituent-dependent modulation of the ^19^F CSA tensor – specifically its magnitude and orientation relative to the C-F bond vector – governs the efficiency of TROSY relaxation interference and can be predictively tuned by chemical design. Guided by these principles, we design a cysteine-reactive 5-fluoropyrimidine reporter, 2,4-dichloro-5-fluoro-5-^13^C-pyrimidine (“2Cl-4Cl”) and experimentally validated its CSA tensor using solid-state MAS NMR. We further demonstrate that solvent-dependent ^19^F and ^13^C chemical shifts define a quantitative environmental susceptibility, enabling calibration of isotropic shifts to local polarity and microenvironment. When installed into a 42-kDa protein, 2Cl-4Cl yields high-resolution dual-nucleus TROSY spectra, substantial reductions in transverse relaxation, and site-specific chemical-shift signatures that distinguish solvent-exposed from partially buried environments. Together, these results transform ^19^F probe development from empirical tagging into a predictive electronic-structure engineering problem, providing a foundation for rational design of relaxation-optimized reporters for large and heterogeneous biomolecular systems.

## RESULTS AND DISCUSSION

### Computational design of relaxation-optimized ^19^F–^13^C spin systems

The broad linewidths of conventional thiol-reactive ^19^F probes arise primarily from their large chemical shift anisotropy (CSA) which, when modulated by rotational tumbling in solution, drives rapid transverse relaxation. Theoretical and experimental studies have shown that aromatic ^19^F-^13^C spin pairs can dramatically narrow these lines when the CSA relaxation mechanism favorably interferes with the dipole-dipole (DD) relaxation mechanism^10^; this phenomenon results in one component of the ^19^F_13C_/^13^C_19F_-coupled doublets relaxing slower than the other (i.e. the transverse relaxation-optimized spectroscopy, or *TROSY*, effect). The efficiency of this interference, however, depends sensitively on the geometry of the CSA tensor relative to the internuclear dipole, rather than on CSA magnitude alone^18^. To quantify this geometric contribution, we define an effective CSA projection,

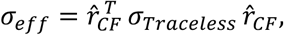

where **σ**_Traceless_ is the traceless CSA tensor of the nucleus of interest and *r̂_CF_* is the unit vector along the C-F bond^18,19^. This quantity represents the component of the CSA tensor that is collinear with the C-F dipolar vector and therefore capable of participating in DD-CSA interference. The sign and magnitude of σ_eff_ therefore determine whether CSA contributes constructively or destructively to transverse relaxation. Importantly, σ_eff_ reflects two distinct physical factors: (i) the overall magnitude of the CSA tensor and (ii) the orientation of that tensor relative to the C-F bond; therefore σ_eff_ sets the maximum possible strength of DD-CSA cross-correlation for a given nucleus. To separate these effects, we additionally consider the normalized quantity |σ_eff_|/Ω, where Ω is the CSA span in Herzfeld-Berger convention (Ω = σ_11_ - σ_33_). This dimensionless ratio isolates tensor geometry from absolute CSA magnitude and provides a measure of dipole-CSA interference efficiency. In intuitive terms, |σ_eff_|/Ω reports how effectively a given CSA tensor is “aimed” along the internuclear dipole, independent of how large the tensor is, which allows scaffolds with very different CSA magnitudes to be compared directly.

As an initial step toward rationally designing a thiol-reactive aromatic scaffold with favorable DD-CSA geometry, we used density functional theory (DFT) to calculate the chemical shielding tensors of three previously reported aromatic ^19^F-^13^C probes: 3-^19^F_13C_-tyrosine, 4-^19^F_13C_-phenylalanine, and 5-^19^F_13C_-uracil (5-FU)^10,20^. Figure 1 depicts the principal axes of the traceless ^19^F-^13^C CSA tensors (Table S1) overlaid on each molecular scaffold, providing a direct geometric visualization of DD-CSA alignment. Using these DFT-derived tensors as inputs (Table S1), we then employed the BRW relaxation theory module of Spinach^21^ to compute the transverse relaxation rates (R_2_) of each ^19^F-^13^C spin pair as a function of magnetic field strength and rotational correlation time (τ_c_) (Fig. 1). Within this theoretical framework, transverse relaxation of heteronuclear single-quantum coherences arises from the combined effects of DD interactions, CSA auto-relaxation, and DD-CSA cross-correlation^22^. While CSA auto-relaxation scales approximately with the square of the CSA magnitude, DD-CSA cross-correlation scales linearly with the projection of the traceless CSA tensor onto the internuclear dipolar frame. All BRW calculations were performed in the rigid limit (S^2^ = 1) assuming an isotropic rotational correlation time (τ_c_) of 25 ns, thereby isolating the influence of CSA geometry alone and providing an upper-bound estimate of transverse relaxation in the absence of internal probe dynamics. Accordingly, these calculations are intended to compare relative behavior across scaffolds, rather than to reproduce experimental R_2_ values quantitatively.

**Figure 1.**
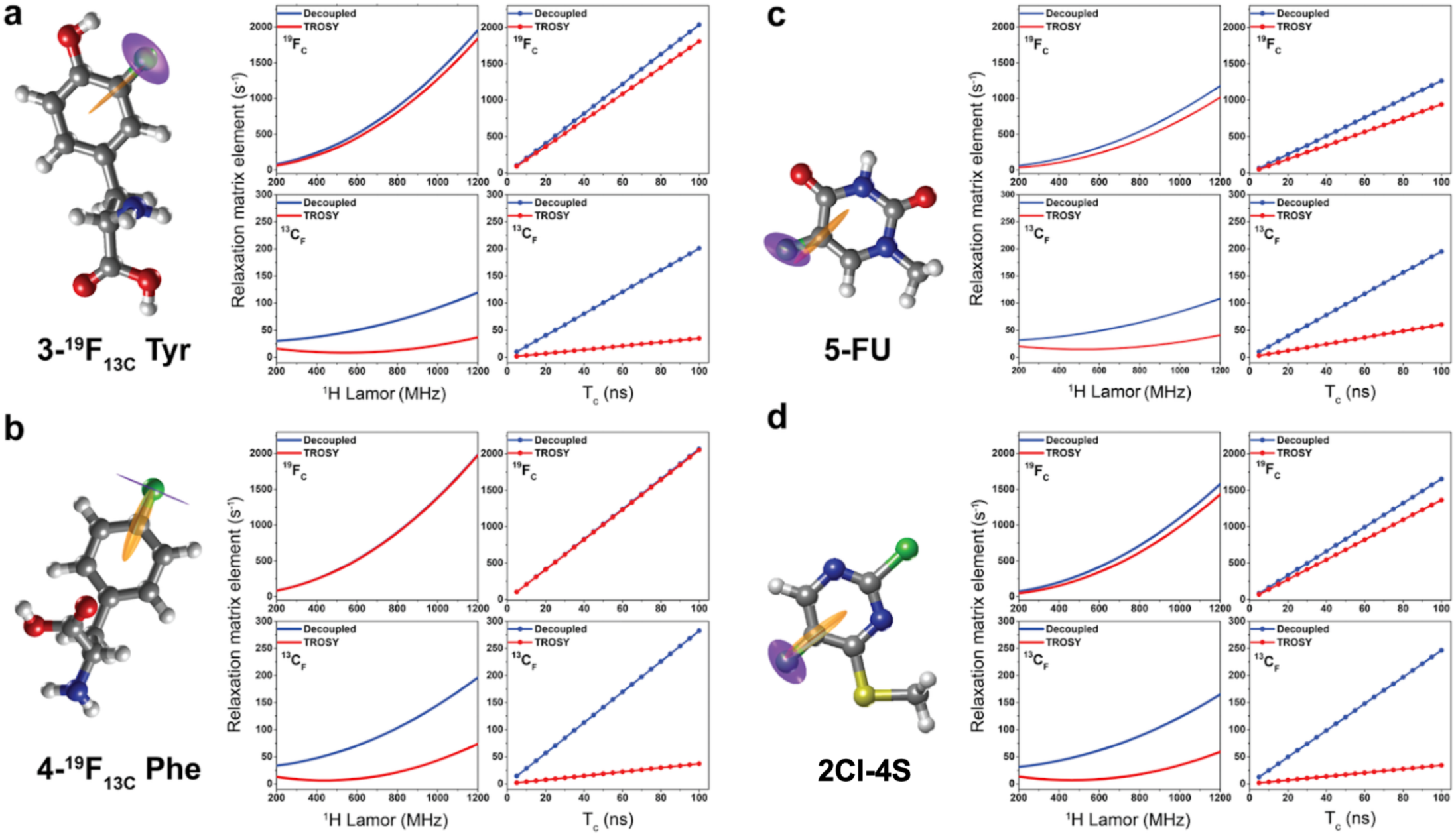
CSA geometry governs DD-CSA interference and ^19^F-^13^C TROSY performance. Principal axes of the traceless ^19^F (purple) and ^13^C (orange) chemical shift anisotropy (CSA) tensors, calculated by density functional theory (DFT), overlaid on the molecular structures of 3-^19^F_13C_-Tyr (**a**) 4-^19^F_13C_-Phe (**b**) 5-^19^F_13C_-uracil (5-FU) (**c**), and 2Cl-4S (**d**). Corresponding Bloch-Redfield-Wangsness (BRW) simulations of TROSY (red) and decoupled (blue) transverse relaxation rates (R_2_) for ^19^F_C_/^13^C_F_ single-quantum coherences are shown as a function of ^1^H Larmor frequency (MHz) and rotational correlation time (τ_c_). All simulations were performed in the rigid limit (S^2^ = 1) assuming isotropic tumbling with τ_c_ = 25 ns, providing an upper-bound, geometry-focused comparison across scaffolds rather than a quantitative reproduction of experimental relaxation rates.

Because 3-^19^F_13C_-Tyr, 4-^19^F_13C_-Phe, and 5-^19^F_13C_-U all exhibit favorable ^13^C transverse relaxation, we focused our design analysis on how the magnitude and orientation of the ^19^F CSA govern probe performance. Examining their chemical shielding tensor properties (Table 1, Table S1), within a framework that considers both the absolute CSA projection (σ_eff_) and its normalization by the CSA span (|σ_eff_|/Ω), clarifies why the phenyl amino acid scaffolds provide such limited ^19^F TROSY benefit. In 3-^19^F_13C_ -Tyr, the predicted ^19^F CSA span is large (Ω ≈ 146 ppm), placing the fluorine nucleus in a regime where CSA auto-relaxation is intrinsically strong. At the same time, one principal component of the ^19^F CSA tensor exhibits substantial alignment with the F-C dipolar axis, giving rise to a non-zero projection (σ_eff_ ≈ −23 ppm; Table 1, Table S1, Fig. 1a). When normalized by its large CSA span, however, this corresponds to a relatively modest interference efficiency (|σ_eff_|/Ω ∼ 0.15), indicating that only a small fraction of the CSA contributes constructively to DD-CSA cross-correlation. As a result, BRW simulations predict that CSA auto-relaxation dominates the relaxation balance over DD-CSA cross-correlation across protein-relevant rotational correlation times (τ_c_ = 10-100 ns; Fig. 1a). In contrast, 4-^19^F_13C_-Phe illustrates the opposite limitation. In this case, the ^19^F CSA principal axis is more closely aligned with the F-C dipolar vector than in 3-^19^F_13C_-Tyr (Fig. 1b, Table S1); however, the associated magnitude is very small, resulting in a near-zero dipolar axis projection (σ_eff_ ≈ 0). Consequently, both σ_eff_ and |σ_eff_|/Ω are negligible, and DD-CSA cross-correlation provides little leverage over relaxation (Table 1, Table S1). Under these conditions, ^19^F transverse relaxation is again dominated by CSA auto-relaxation across the full range of τ_c_ examined (Fig. 1b). Within this design framework, these two phenyl amino acids therefore illustrate distinct failure modes: poor ^19^F relaxation can arise either from excessive CSA magnitude despite reasonable geometric alignment (3-^19^F_13C_-Tyr), or from insufficient CSA projection even with very favorable orientation (4-^19^F_13C_-Phe). In 5-^19^F_13C_-U, both limitations are alleviated within the same theoretical analysis. The CSA span is substantially smaller than those of phenyl-based amino-acid scaffolds (Ω ≈ 110 ppm versus 142-156 ppm), reducing the CSA auto-relaxation penalty (Table 1, Table S1). At the same time, the rhombic character of the ^19^F CSA tensor produces a meaningful dipolar-axis projection (σ_eff_ ≈ −32 ppm), yielding a substantially larger normalized interference efficiency (|σ_eff_|/Ω ∼ 0.3). This combination of reduced CSA magnitude and enhanced geometric efficiency enables effective DD-CSA cross-correlation without incurring a prohibitive auto-relaxation penalty. Accordingly, BRW simulations predict balanced DD-CSA interference and substantially improved ^19^F transverse relaxation (Fig. 1c).

**Table 1.** DFT-derived CSA tensor parameters defining DD-CSA interference in aromatic ^19^F-^13^C probes.

| Molecule | Span $\Omega$<br>(ppm) | Skew<br>$\kappa$ | $\sigma_{\text{eff}}$<br>(ppm) | $\vert \sigma_{\text{eff}}\vert /\Omega$ |
| --- | --- | --- | --- | --- |
| **3-$^{19}\text{F}_{13\text{C}}$-Tyr** |  |  |  |  |
| $^{19}\text{F}$ | 146.30 | -0.47 | -23.40 | 0.16 |
| $^{13}\text{C}$ | 124.15 | 0.09 | -63.15 | 0.51 |
| **4-$^{19}\text{F}_{13\text{C}}$-Phe** |  |  |  |  |
| $^{19}\text{F}$ | 152.58 | 0.04 | 1.76 | 0.01 |
| $^{13}\text{C}$ | 164.88 | 0.30 | -90.66 | 0.55 |
| **5-$^{19}\text{F}_{13\text{C}}$-U** |  |  |  |  |
| $^{19}\text{F}$ | 109.78 | -0.64 | -32.06 | 0.29 |
| $^{13}\text{C}$ | 114.38 | 0.19 | -49.52 | 0.43 |
| **2Cl-4S** |  |  |  |  |
| $^{19}\text{F}$ | 126.15 | -0.69 | -29.28 | 0.23 |
| $^{13}\text{C}$ | 149.67 | 0.28 | -80.47 | 0.54 |

These results made 5-fluorouracil (5-^19^F_13C_-U), which was previously used for TROSY-NMR studies of RNA by Becette et al^23^, a logical starting point for designing a thiol-reactive ^19^F-^13^C TROSY-optimized probe. Our goal was not necessarily to outperform 5-FU in absolute ^19^F relaxation, but to preserve its favorable CSA geometry while enabling site-selective thiol reactivity. While 5-fluorouracil itself is not reactive toward nucleophiles, a standard thiol reactive functional group, such as bromoacetamide or acrylamide or thiosulfonate (maleimide is unsuitable since it introduces two diastereomeric products) can in principle be added via a linker attached to the uracil ring. However, this requires protection of one of the two nucleophilic nitrogens in the pyrimidine ring, in order to allow regioselective derivatization of one nitrogen, and to avoid potential reactions with the electrophilic “warhead”. Alternatively, we wondered whether 5-FU can be modified so as to allow direct addition of the cysteine sulfhydryl group, e.g. via aromatic nucleophilic substitution (S_N_Ar) at position C4 (Scheme 1). A search of the literature revealed that a dichlorinated derivative of 5-FU (2,4-dichloro-5-fluoropyrimidine, “2Cl-4Cl”, Scheme 1) reacts readily with various nucleophiles, including thiols, and is easily accessible by treatment of 5-FU with POCl_3_^24^. The mono-chlorinated 5-FU analog (2O-4Cl, Scheme 1) is very sparsely reported in the literature, but appears to be less electrophilic, and its synthesis from 5-FU via a thioxo derivative is not as efficient as that of the dichloro derivative^24,25^. Moreover, 4-chloro-5-fluorouracil (2O-4Cl) can exist in three possible tautomeric forms, which, as suggested recently^26^ may enhance its fluorine chemical shift dispersion, but would complicate our theoretical approach.

**Scheme 1.**
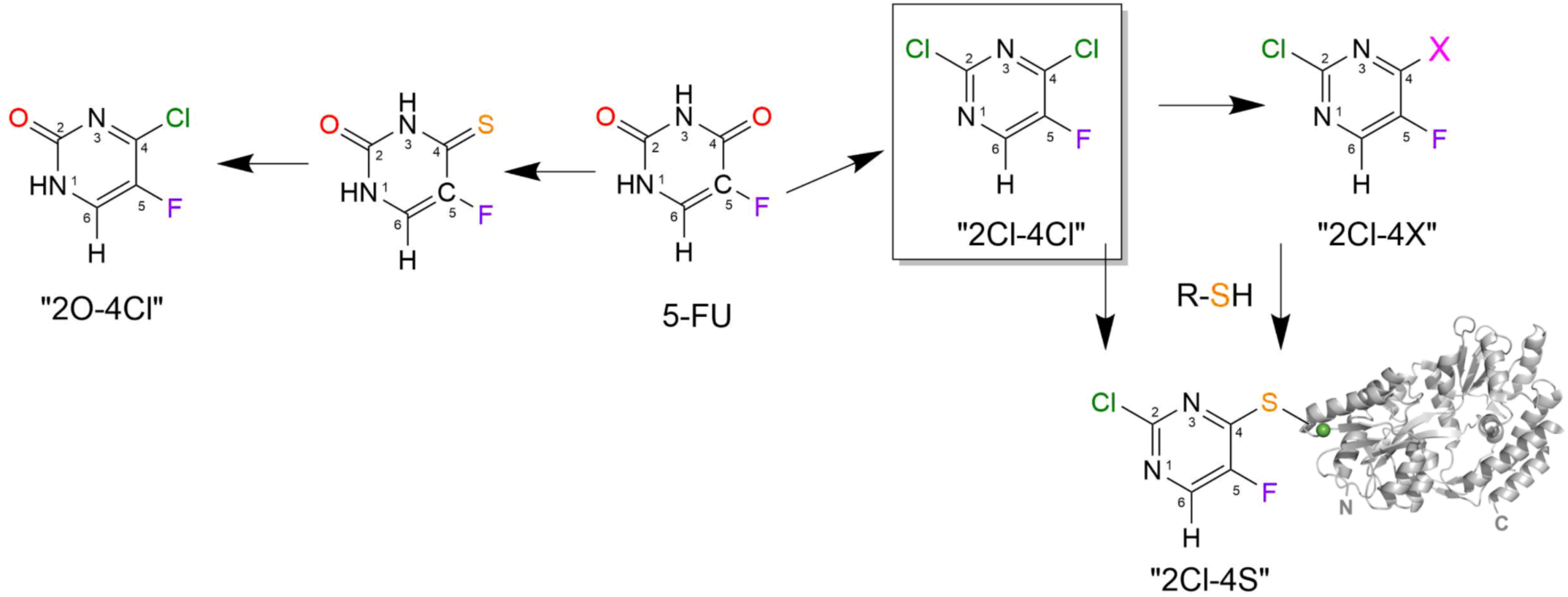
Modifications of 5-fluorouracil to introduce thiol reactivity in the pyrimidine ring.

To evaluate whether the 2-chloro analogs of 5-FU retain its key DD-CSA design features, we computed the chemical shielding tensors for both 2,4-dichloro-5-fluoropyrimidine (“2Cl-4Cl”, Scheme 1) and its thiol-substituted product (“2Cl-4S”), and analyzed their σ_eff_ and |σ_eff_|/Ω (Fig. S1, Table S1, and Table 1). For ^19^F, both chlorinated scaffolds maintain substantial dipolar-axis projections (2Cl-4Cl: σ_eff_ ≈ −32 ppm, |σ_eff_|/Ω ∼ 0.24; 2Cl-4S: σ_eff_ ≈ −29 ppm, |σ_eff_|/Ω ∼ 0.23), indicating that the geometry required for DD-CSA interference is preserved. Nonetheless, rigid-limit BRW simulations show that 5-FU remains the most favorable scaffold for absolute ^19^F transverse relaxation in this set (Fig. 1): 5-FU yields the lowest predicted ^19^F TROSY and decoupled R_2_ values (229.9 s^−1^ and 315.5 s^−1^, respectively), whereas 2Cl-4S and the unreacted chlorinated precursor exhibit higher ^19^F R_2_ values (e.g., 2Cl-4S: 337.4 s^−1^ TROSY; 414.0 s^−1^ decoupled). Thus, chlorination and thiol substitution do not improve the absolute ^19^F relaxation baseline under rigid-limit conditions, even though they retain meaningful interference efficiency. However, the same modifications are predicted to strongly enhance ^13^C performance. Both chlorinated scaffolds exhibit large ^13^C dipolar-axis projections and high normalized efficiencies (2,4-dichloro-5-fluoropyrimidine: σ_eff_ ≈ −74 ppm, |σ_eff_|/Ω ∼ 0.5; 2Cl-4S: σ_eff_ ≈ −80 ppm, |σ_eff_|/Ω ∼ 0.5) consistent with particularly favorable CSA alignment along the C-F bond (Table 1, Table S1). Correspondingly, BRW simulations predict substantially slower ^13^C TROSY relaxation for both 2Cl-4S and the unreacted scaffold 2Cl-4Cl compared to 5-FU (e.g., 4Cl-2Cl: 5.4 s^−1^ versus 5-FU: 13.1 s^−1^), while maintaining robust TROSY/anti-TROSY separation (Fig. 1, Fig. S1).

Taken together, 4Cl-2Cl represents a deliberately engineered compromise that preserves 5-FU-like interference geometry while potentially enabling thiol-based incorporation and enhanced ^13^C TROSY performance, at the cost of increased absolute ^19^F relaxation.

### Thiol-reactivity of 2Cl-4Cl in aqueous buffer

The reaction of 2Cl-4Cl with glutathione in aqueous buffer was monitored by NMR (Figure 2) and ESI-MS. Reaction between the natural abundance compound (2 mM) and a slight excess of glutathione in phosphate buffer at pH 7.2 was complete in 3-5 hours, giving a single product despite the presence of five-fold excess of lysine over glutathionine. ESI-MS showed the disappearance of the GSH signal and the appearance of a GSH-2Cl-4S adduct with concomitant loss of chloride. The product was stable at room temperature in aqueous buffer for at least one month (**Figs. S2, S3**).

**Figure 2.**
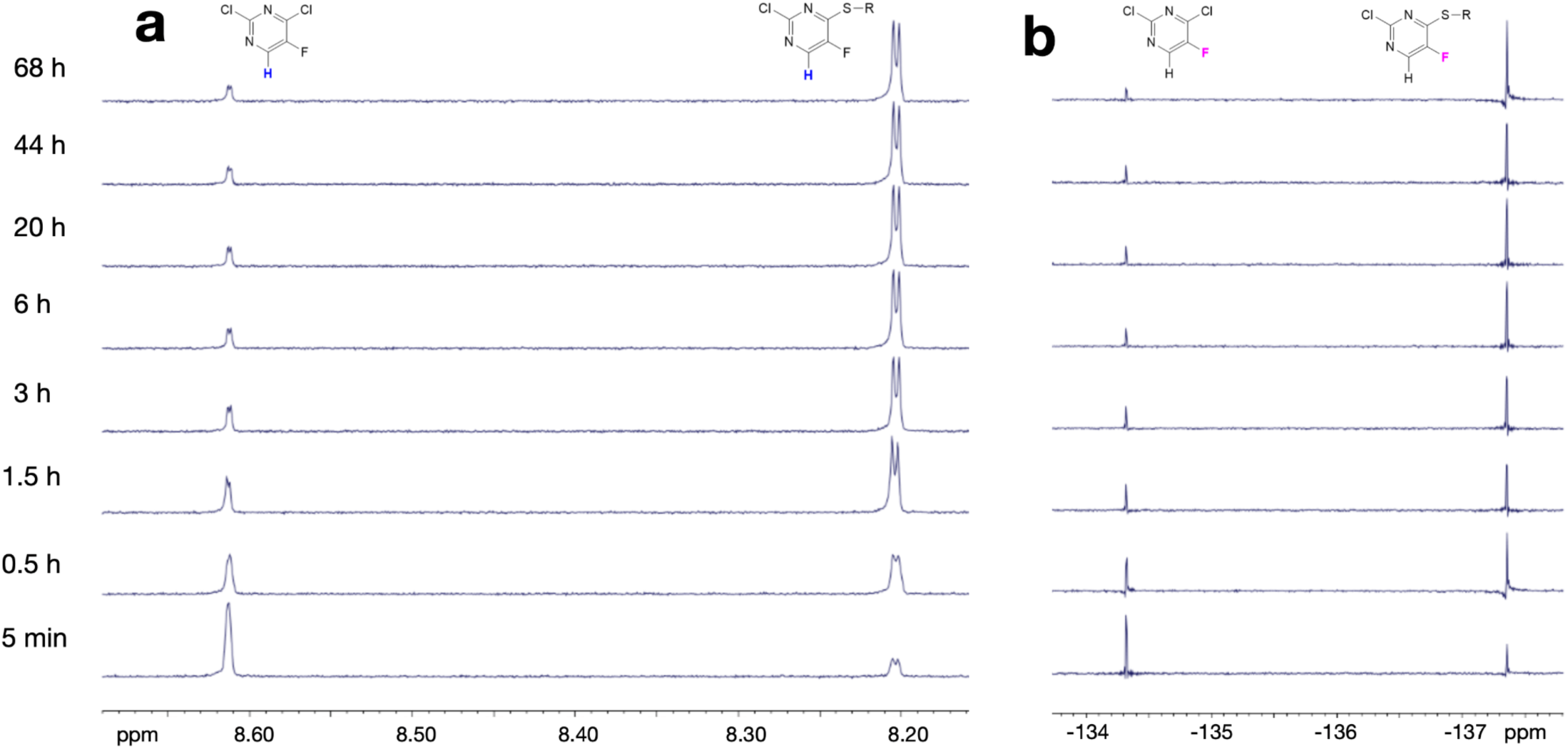
Thiol-specific reactivity of 2Cl-4Cl in dilute aqueous solution containing excess lysine. ^1^H-NMR (**a**) and ^19^F-NMR (**b**) spectra of 2Cl-4Cl (natural abundance, 2 mM) in the presence of 2.5 mM glutathione and 10 mM lysine in 50 mM sodium phosphate D_2_O buffer at pH 7.2.

**Scheme 2.**
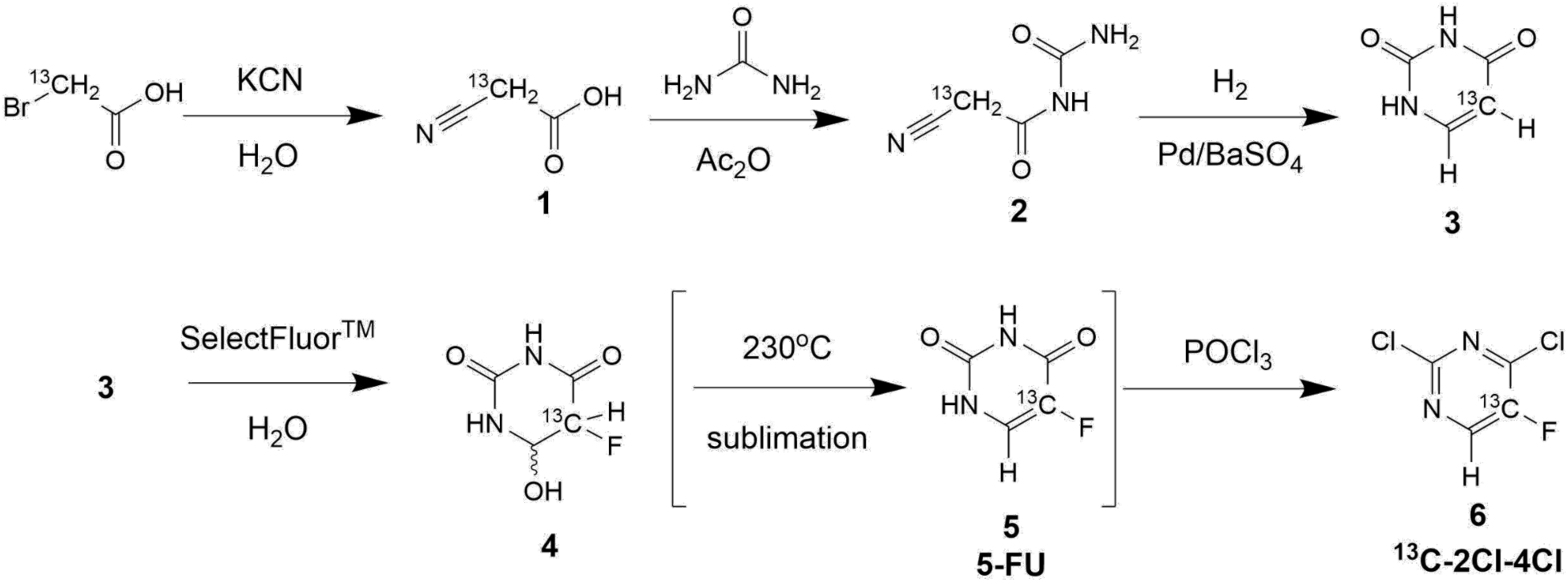
Synthesis of the ^13^C-labeled 2,4-dichloro-5-fluoropyrimidine (“2Cl-4Cl”).

Given that many biologically interesting proteins are less soluble than GSH, we attempted to enhance the thiol-reactivity of this probe by replacing the chloride with a better leaving group. The order of leaving group reactivity in S_N_Ar reactions can be difficult to anticipate^27^, but generally follows the order of electronegativity, rather than anion stability (e.g. F > Cl ∼ Br > I). The 4-fluoro-substituted analog (2Cl-4F, Scheme 1) was therefore expected to be more active and was synthesized by treating 2Cl-4Cl with CsF in DMSO. This derivative was indeed an order of magnitude more reactive toward glutathione, but was prone to competing hydrolysis and lysine reactions (data not shown). In an attempt to fine-tune the reactivity of the probe further, we attempted to replace the chloride at position 4 with a triflate, cyanide, and nitro- groups. We could not obtain the nitro- and triflate-substituted analogs, and the 4-cyano analog (2Cl-4CN) was found to be unreactive toward GSH (data not shown). Based on these results, 2Cl-4Cl was chosen as the most suitable 5-FU analog for further studies, as its somewhat sluggish reactivity toward thiols is compensated by an impressive inertness toward lysine and water. The 0.5-1.0 M stock solutions in DMSO are suitable for cysteine labeling of proteins (*vide infra*) and are stable for several weeks if stored in the freezer.

### Synthesis of ^13^C-labeled 2Cl-4Cl

The synthesis of 5-^13^C-labeled 2Cl-4Cl is shown in Scheme 2. ^13^C-labeled uracil (**3**) was synthesized following the well-established protocols^28–30^. Briefly, reaction of ^13^C-bromoacetic acid with cyanide yielded cyanoacetic acid (**1**), which was condensed with urea and cyclized under reductive conditions to give 5-^13^C-uracil (**3**) in 69% overall yield. Fluorination of the labeled uracil (**3**) with SelectFluor^TM31^ gave 5-fluoro-hydroxydihydrouracil (**4**) which, when heated under vacuum and sublimated to give fluorouracil (**5**). The chlorinated derivative (2Cl-4Cl) can be obtained from 5-FU by treatment with POCl_3_^32^. Skipping the fluorouracil isolation, the crude fluorination product (**4**) was simultaneously dehydrated and chlorinated with POCl_3_, giving ^13^C-labeled 2Cl-4Cl in 21% overall yield from ^13^C-bromoacetic acid.

### Experimental CSA tensors validate the DD-CSA design rules

To experimentally test the predictions of the design framework described above, we determined the full ^19^F chemical-shift anisotropy tensors of 5-FU and 2,4-dichloro-5-fluoropyrimidine using solid-state magic-angle spinning (MAS) NMR. Measurements were performed on powdered samples under identical conditions to ensure direct comparability, with ^19^F{^1^H} MAS spectra acquired at two spinning frequencies to verify robustness of the extracted tensor parameters (Figure 3a). The 2,4-dichloro-5-fluoropyrimidine spectrum exhibits a well-defined spinning sideband manifold that is accurately reproduced by a single CSA tensor, whereas the 5-FU spectrum requires two closely related CSA tensor components to reproduce its spinning sideband manifold, indicating the presence of multiple inequivalent fluorine environments in the solid state (Figure 3a). Each 5-FU tensor component exhibits increased rhombicity relative to 2,4-dichloro-5-fluoropyrimidine, consistent with reduced local symmetry, but the multiplicity itself most likely reflects distinct packing or intermolecular interaction motifs rather than CSA shape alone. Quantitative fitting of the MAS line shapes was performed using DMFit^33^ and the resulting CSA tensors are reported in the Herzfeld-Berger convention in terms of the span (Ω) and skew (κ).

**Figure 3.**
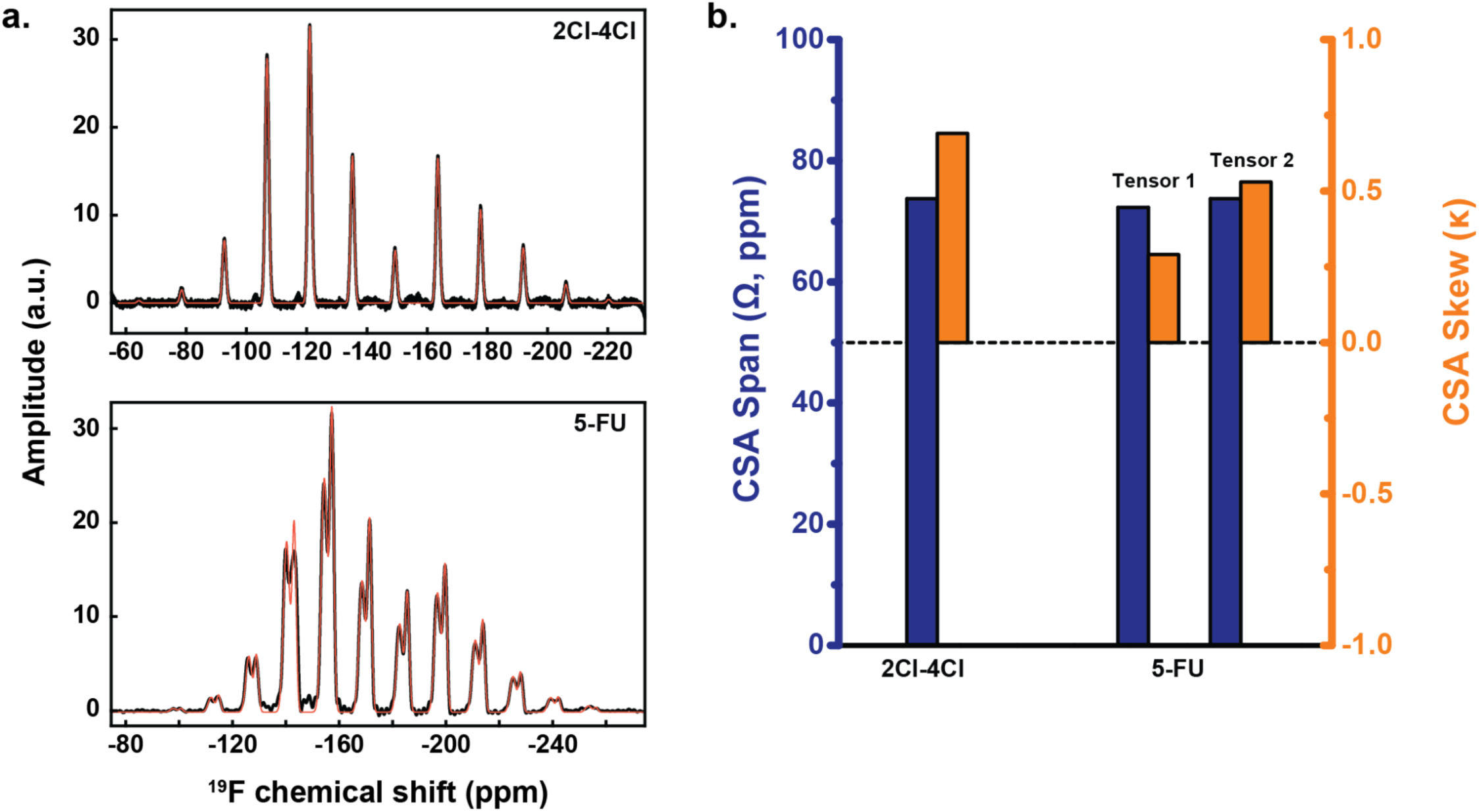
Experimental validation of CSA design principles. **a**, ^19^F MAS spectra of 2,4-dichloro-5-fluoropyrimidine and 5-FU at two spinning frequencies (8 kHz and 30 kHz; black) overlaid with DMFit simulations (orange). **b**, DMFit was used to extract CSA span (Ω) and skew (κ) in the Herzfeld-Berger convention for 2,4-dichloro-5-fluoropyrimidine and 5-FU.

For 2,4-dichloro-5-fluoropyrimidine, the extracted CSA tensor is characterized by a span Ω = 73.7 ppm and a skew κ = 0.69, indicating a pronounced axial bias of the shielding tensor (Fig. 3b). 5-FU exhibits comparable CSA spans (Ω ≈ 72.3 and 73.7 ppm) but substantially lower skew values (κ = 0.29 and 0.53), reflecting a more rhombic tensor shape (Fig. 3b). Importantly, although the two compounds display similar CSA magnitudes, the increased axial character of the 2,4-dichloro-5-fluoropyrimidine tensor is expected to reduce the CSA auto-correlation penalty and favor more effective destructive interference between CSA- and dipolar-mediated relaxation pathways. The experimentally determined CSA tensors closely match the tensor-shape trends predicted by DFT calculations (Fig. 1, Table 1), which indicated that chlorine substitution at the 4-position enhances axial bias of the ^19^F shielding tensor and alters its orientation relative to the molecular frame. While these solid-state MAS measurements do not directly determine the absolute orientation of the CSA principal axes, they nonetheless establish that substituent-controlled modulation of CSA symmetry constitutes a physically realizable and quantitatively-validated mechanism for reducing transverse relaxation in aromatic ^19^F probes.

### 2Cl-4S demonstrates simultaneous ^19^F and ^13^C TROSY optimization in proteins

Having shown that CSA geometry can be deliberately tuned at the molecular level, we next asked whether this design strategy remains effective after incorporation into a protein. To address this question, ^13^C-2Cl-4Cl was site-specifically conjugated to maltose-binding protein (MBP), a soluble 42-kDa system whose global rotational correlation time is estimated to be ∼ 25 ns^10^. We selected two solvent-exposed residues, K34 and R354, and generated corresponding single-and double-cysteine mutants (K34C and K34C/R354C). These positions were chosen based on their spatial separation and putative minimal structural impact (Fig. 4a). Labeling of engineered cysteine sites proceeded cleanly under protein-compatible aqueous conditions, reacting to completion with ∼ 6h, and yielding single fluorinated adducts as confirmed by intact mass spectrometry (Figs. S4). 1D ^19^F NMR spectra collected at 600 MHz revealed that each labeling site produced a distinct pair of TROSY and anti-TROSY resonances (Fig. 4b). Each doublet collapsed into a singlet upon ^13^C decoupling, confirming the presence of a one-bond scalar coupling to ^13^C (¹*J*_CF_ ≈ 265 Hz), which lies at the upper end of the typical range reported for aromatic ^19^F-^13^C pairs. High-resolution 2D ^19^F-^13^C TROSY-HSQC spectra acquired using an “out-and-stay” style pulse scheme^10^ contained two well-resolved cross peaks (Fig. 4c).

**Fig. 4.**
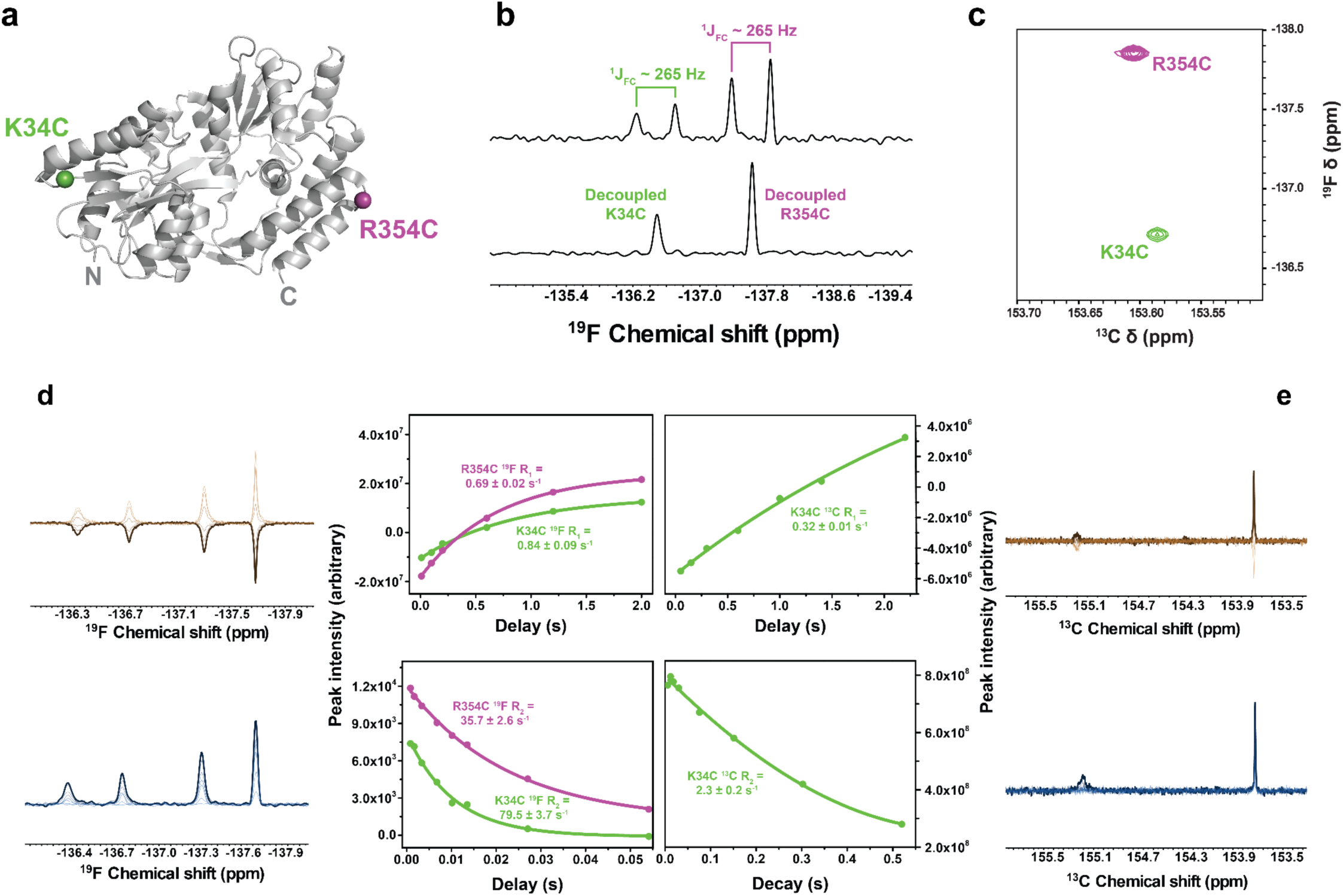
Experimental validation of CSA-geometry-guided TROSY optimization in a protein. **a,** Structural model of maltose-binding protein (MBP, 42 kDa; PDB 8C8F) highlighting solvent-exposed labeling sites K34 and R354 used for site-specific conjugation of 2Cl-4Cl. **b,** 1D ^19^F NMR spectra (600 MHz) of MBP labeled at K34C and R354C, showing resolved TROSY and anti-TROSY doublets arising from ¹*J*_CF_ ≈ 265 Hz (*top*); spectra collapse to singlets upon ^13^C decoupling (*bottom*). **c,** 2D ^19^F-^13^C TROSY-HSQC spectrum acquired using an out-and-stay pulse scheme, revealing two well-resolved cross-peaks assigned to K34C and R354C from the corresponding 1D ^19^F spectrum. **d,** Site-dependent transverse (R_2_) and longitudinal (R_1_) relaxation rates for the ^19^F TROSY and anti-TROSY components, quantified from one-dimensional relaxation experiments. **e,** ^13^C transverse (R_2_) and longitudinal (R_1_) relaxation rates at the K34C site.

We next quantified transverse relaxation rates for both TROSY and anti-TROSY components using 1D NMR dynamics experiments; in general, the measured R_2_ rates of the TROSY components are reduced by approximately twofold relative to the anti-TROSY components. The ^19^F TROSY component of 2Cl-4S-K34C exhibits an R_2_ of 79.5 ± 3.7 s^−1^, while at R354C the corresponding ^19^F TROSY R_2_ is reduced to 35.7 ± 2.6 s^−1^ (Fig. 4d; Table 2). In the carbon dimension, transverse relaxation is substantially slower overall with a ^13^C TROSY R_2_ value of 2.28 ± 0.2 s^−1^ at K34C (Fig. 4e); we did not quantify ^13^C relaxation at R354C owing to spectral overlap in the double-mutant (Fig. S4). In contrast, longitudinal relaxation (R_1_) rates for both nuclei vary only modestly between TROSY and anti-TROSY coherence pathways (Table 2), indicating that the observed gains arise specifically from suppression of transverse relaxation rather than changes in global tumbling or probe mobility.

**Table 2.**
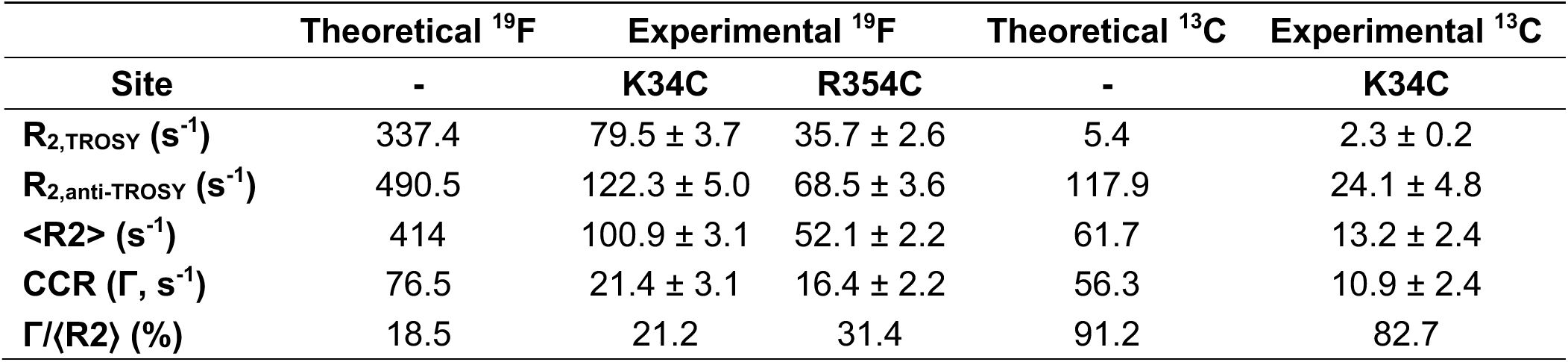
Experimental transverse relaxation (R2) and DD-CSA cross-correlation (Γ) rates for 2Cl-4Slabeled MBP.

To facilitate mechanistic interpretation, we define the component-averaged transverse relaxation rate (⟨R_2_⟩; ⟨R_2_⟩ = [R_2,TROSY_ + R_2,anti_]/2). Physically, ⟨R_2_⟩ reflects the overall strength of transverse relaxation experienced by the coupled spin pair, independent of how that relaxation is partitioned between the two coherence pathways. Because both TROSY and anti-TROSY R_2_ values were measured directly, the experimental dipole-CSA cross-correlated relaxation (CCR; Γ) rates could be extracted from their differences (Γ = [R_2,anti_ - R_2,TROSY_]/2)^18^. The resulting CCR terms account for a substantial fraction of ⟨R_2_⟩, directly demonstrating the presence of DD-CSA interference in the protein environment. For ^19^F, Γ values of 21.4 ± 3.1 s^−1^ at K34C and 16.4 ± 2.2 s^−1^ at R354C correspond to ∼21% and ∼31% of ⟨R_2_⟩, respectively (Table 2). A similarly pronounced effect is observed in the carbon dimension, where the ^13^C CCR at K34C accounts for ∼83% of ⟨R_2_⟩, indicating that DD-CSA interference dominates the transverse relaxation behavior of the carbon coherence (Table 2). This Γ/⟨R_2_⟩ fractional representation provides a direct bridge between experiment and theory. In the design framework developed above, the dimensionless quantity |σ_eff_|/Ω describes how efficiently the CSA tensor projects onto the C-F dipolar vector, independent of its absolute magnitude. Experimentally, the ratio Γ/⟨R_2_⟩ plays an analogous role: it reports what fraction of the observed transverse relaxation is governed by dipole-CSA interference rather than by non-interfering relaxation processes. Consistent with this interpretation, the theoretical Γ/⟨R_2_⟩ values are comparable in magnitude to those observed experimentally, despite substantially overestimating absolute relaxation rates (Table 2). This agreement at the level of dimensionless ratios indicates that while local probe dynamics strongly attenuate overall relaxation rates in proteins, they do not eliminate the underlying geometric interference encoded by the CSA tensor. At the K34C site, both Γ and ⟨R_2_⟩ are reduced by similar factors relative to rigid-limit predictions (Table 2), indicating that local probe dynamics largely rescale transverse relaxation without altering the relative contribution of dipole-CSA interference. At the R354C site, ⟨R_2_⟩ is more strongly suppressed than Γ, resulting in an increased Γ/⟨R_2_⟩ ratio (Table 2); this site dependence is most naturally attributed to differences in local probe mobility rather than changes in CSA-dipole geometry, which is fixed by the molecular scaffold.

Together, these results show that the fraction of transverse relaxation arising from DD-CSA interference, measured experimentally as Γ/⟨R_2_⟩, provides a direct and experimentally accessible analogue of the theoretical geometry metric |σ_eff_|/Ω, validating CSA geometry as a robust and transferable design parameter for simultaneous ^19^F and ^13^C TROSY optimization in proteins.

### Effects of slow tumbling on ^19^F and ^13^C transverse relaxation

Having demonstrated that 2Cl-4S enables simultaneous ^19^F and ^13^C TROSY optimization in MBP, we next examined how this behavior evolves as molecular tumbling slows into regimes characteristic of larger macromolecular assemblies. Because transverse relaxation arising from DD and CSA interactions depends strongly and nonlinearly on the rotational correlation time (τ_c_), it is essential to evaluate both the robustness and the practical limits of the CSA-geometry design strategy under slow-tumbling conditions. To access a controlled range of correlation times, 2Cl-4S-labeled MBP(K34C/R354C) was studied at increasing solvent viscosity achieved through combined temperature reduction and glycerol addition. Dynamic viscosities were calculated using the Cheng correlation^24^, and corresponding correlation times were estimated by scaling from a reference τ_c_ of ∼25 ns for MBP in aqueous buffer at 25 °C using standard Stokes-Einstein-Debye relationships (Fig. 5a). Across this range, 2D out-and-stay TROSY-HSQC spectra were acquired using both ^19^F detection (i.e. ^13^C-^19^F) and ^13^C detection (i.e. ^19^F-^13^C) modes, enabling a direct comparison of how slow tumbling differentially impacts the two nuclei (Fig. 5b,c).

**Fig. 5.**
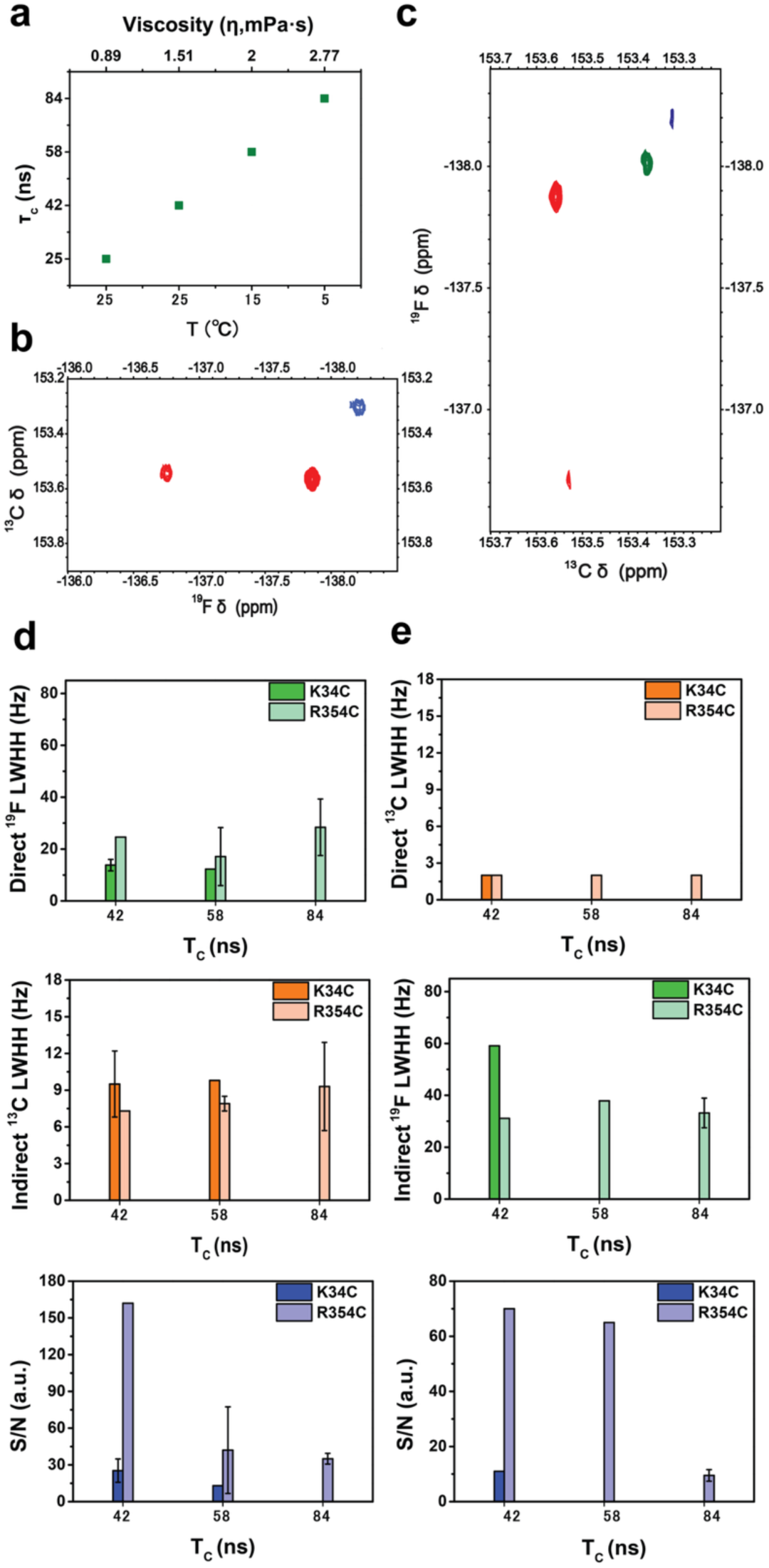
Differential ^19^F and ^13^C relaxation behavior under slow tumbling. **a,** Dynamic viscosities of 20% (w/v) glycerol solution were calculated using the Cheng correlation^24^, and corresponding rotational correlation times (τ_c_) were estimated by scaling from a reference τ_c_ ∼ 25 ns for MBP in aqueous buffer at 25 °C using Stokes-Einstein-Debye relationships. Representative (**b**) ^13^C-^19^F and (**c**) ^19^F-^13^C “out-and-stay” TROSY spectra acquired under different viscosities to simulate ∼42 ns (∼121 kDa, red), ∼56 ns (∼156 kDa, green), and ∼78 ns (∼230 kDa, blue) conditions. **d,** Summary of linewidths at half-height (LWHH) for 2Cl-4S-MBP(K34C/R354C) resonances in ^13^C-^19^F TROSY spectra for ^19^F (directly-acquired, top) and ^13^C (indirectly-acquired, middle) as a function of temperature, with corresponding τ_c_ values indicated, together with signal-to-noise (S/N) ratios (bottom). **e,** Summary of linewidths at half-height (LWHH) for 2Cl-4S-MBP(K34C/R354C) resonances in ^19^F-^13^C TROSY spectra for ^13^C (directly-acquired, top) and ^19^F (indirectly-acquired, middle) as a function of temperature, with corresponding τ_c_ values indicated, together with signal-to-noise (S/N) ratios (bottom). The digital resolution in the directly-detected dimension of all spectra exceeded the measured linewidths by at least a factor of two.

As τ_c_ increased from ∼42 ns to ∼78 ns, corresponding to effective molecular weights of ∼120-230 kDa, the two detection modes exhibit qualitatively distinct responses. In these experiments, the apparent linewidth at half height (LWHH) scales directly with the transverse relaxation rate (R_2_); thus, changes in LWHH directly report changes in transverse relaxation rather than resolution or processing artifacts. The apparent ^13^C TROSY linewidths in the carbon-detected experiments remain essentially unchanged across the entire τ_c_ range examined, remaining narrow even at the highest effective molecular weights (Fig. 5d). This behavior reflects the persistence of favorable DD-CSA cross-correlation for the ^13^C TROSY coherence. Although slow tumbling is predicted to rapidly suppresses the anti-TROSY component (Fig. 1), the TROSY-selected ^13^C coherence remains protected against additional transverse relaxation. Once this slow-tumbling regime is reached, further increases in τ_c_ do not substantially shorten the ^13^C transverse coherence lifetime, resulting in an effective linewidth plateau. The experimentally observed linewidth trends mirror the qualitative behavior predicted by BRW calculations. In the theoretical τ_c_-dependent R_2_ profiles, the ^13^C TROSY component exhibits a markedly weaker dependence on the rotational correlation time than either the ^19^F TROSY or anti-TROSY components, while TROSY/anti-TROSY separation persists into the slow-tumbling regime (Fig. 1d). Although the absolute experimental linewidths are substantially smaller due to local probe dynamics and motional averaging not captured in the rigid-limit calculations, the agreement at the level of τ_c_-dependent trends supports the interpretation that the observed linewidth plateau arises from the same DD-CSA interference mechanisms identified theoretically.

In contrast, fluorine-detected spectra show a stronger dependence on τ_c_. Apparent ^19^F linewidths broaden with increasing viscosity, reflecting the combined effects of CSA auto-relaxation and the heightened sensitivity of ^19^F chemical shifts to conformational exchange (Fig. 5e). Importantly, this broadening does not simply erode spectral information. At lower temperatures, several ^19^F resonances undergo clear spectral splitting rather than unresolved line broadening, and these splittings persist under proton decoupling during acquisition (Fig. 5c). This behavior is consistent with the emergence of distinct, slowly interconverting conformational or microenvironmental substates that become spectroscopically resolvable as exchange slows^34^ highlighting the sensitivity of fluorine detection to heterogeneity under slow-tumbling conditions.

Despite the near-constant ^13^C linewidths, the signal-to-noise ratio (S/N) of the carbon-detected spectra decreases substantially with increasing τ_c_ (Fig. 5b,5d). This apparent decoupling of linewidth and sensitivity reflects a fundamental distinction between transverse relaxation and signal intensity. Linewidth reports the lifetime of an individual coherence once formed, whereas S/N depends on how efficiently such coherences are generated and detected. In the slow-tumbling regime, longitudinal relaxation (R_1_) slows, leading to incomplete recovery of ^13^C magnetization between scans and increased saturation under fixed recycle delays^34^ In addition, although carbon detection avoids the most severe effects of rapid ^19^F transverse relaxation during acquisition, coherence transfer from ^19^F to ^13^C remains gated by fluorine relaxation; as τ_c_ increases, a progressively smaller fraction of spins survives this transfer step. These effects reduce the total observable signal without further broadening the linewidth of the surviving ^13^C TROSY coherence. Importantly, in all experiments the digital resolution in the directly-detected dimension exceeded the measured linewidths by at least a factor of two, confirming that the reported linewidths reflect intrinsic transverse relaxation rather than instrumental or processing limitations. Together, these observations show that under slow-tumbling conditions ^13^C detection preserves high spectral resolution but becomes sensitivity-limited, whereas ^19^F detection increasingly emphasizes conformational heterogeneity and exchange. The complementary behavior of the two nuclei defines both the strengths and the intrinsic trade-offs of simultaneous ^19^F/^13^C TROSY optimization in large macromolecular systems.

## CONCLUSION

Chemical shift anisotropy has long been recognized as the dominant relaxation mechanism governing the performance of aromatic ^19^F probes in biomolecular NMR, yet it has typically been treated as an immutable nuclear property rather than a chemically tunable variable. In this work, we demonstrate that the geometry of the CSA tensor, specifically its orientation and projection relative to the internuclear dipolar interaction, constitutes a decisive and engineerable determinant of transverse relaxation. By reframing CSA as a vectorial quantity whose alignment can be predictively controlled through molecular design, we establish a general framework for rationally shaping dipole-CSA interference in coupled ^19^F-^13^C spin systems, extending foundational concepts of cross-correlated relaxation into a chemical design paradigm^18,35^ Beyond one-dimensional readouts, the engineered ^19^F-^13^C spin system enables multidimensional correlation experiments and provides a practical platform for multidimensional relaxation and exchange measurements in site-specifically labeled proteins.

Through an integrated combination of electronic-structure calculations, Bloch-Redfield-Wangsness relaxation analysis, solid-state MAS NMR, and protein-based validation, we show that substituent-dependent modulation of CSA tensor symmetry and orientation enables deliberate control over relaxation pathways. Experimental determination of CSA tensors confirms the predicted trends in tensor shape and validates the physical basis of the design strategy. When implemented in a cysteine-reactive fluoropyrimidine scaffold, these principles yield a probe that supports simultaneous ^19^F and ^13^C TROSY optimization in a 42 kDa protein, with DD-CSA cross-correlated relaxation accounting for a substantial fraction of the observed transverse relaxation. This agreement between theoretical geometry metrics and experimentally measured cross-correlated relaxation demonstrates that CSA geometry is a robust and transferable design parameter that survives incorporation into a protein environment.

Extension of these measurements into the slow-tumbling regime establishes that the most consequential dynamical implication of the 2Cl-4S scaffold is the exceptionally slow transverse relaxation of the directly bonded ^13^C nucleus. Carbon-detected TROSY coherences remain protected against additional transverse relaxation even as effective molecular weights exceed 200 kDa, consistent with sustained CSA-dipolar interference and resulting in linewidth plateaus rather than progressive broadening. From a first-principles perspective, a baseline ^13^C transverse relaxation rate on the order of 2-3 s^−1^ places this nucleus in an unusually favorable regime for relaxation-dispersion experiments: long transverse coherence lifetimes permit extended relaxation delays, minimize systematic inflation of baseline R_2_, and ensure that even modest exchange-induced contributions would represent a significant and readily quantifiable perturbation^34,36^.

More broadly, the framework established here transforms relaxation optimization from an empirical outcome into a chemically addressable design problem. Because CSA geometry is dictated by local electronic structure, these concepts are not specific to a single scaffold but are readily extensible to other aromatic systems, substituent patterns, and heteronuclear spin pairs. By placing relaxation control on the same conceptual footing as reactivity and selectivity, this work provides a foundation for the rational development of next-generation NMR probes tailored to the size, dynamics, and heterogeneity of increasingly complex biomolecular systems.

## METHODS

### DFT relaxation calculations

Geometries of all molecules were optimized in *Gaussian16*^37^using M06 exchange-correlation functional^38^ with cc-pVDZ basis set ^39–41^. In structures that contained iodine, LANL2DZdp ECP basis set^42–44^ was used for iodine atoms. Chemical shielding tensors were computed using GIAO method^45^ Detailed Gaussian calculation logs are included into the Supplementary Information.

Atomic coordinates and chemical shielding tensors of the active ^13^C-^19^F pair in each molecule were imported into *Spinach* 2.10^21^, where Redfield relaxation superoperators were calculated numerically (as described in^46^) using isotropic rotational diffusion approximation with a rotational correlation time of 25 ns. Transverse relaxation rates of TROSY and anti-TROSY components of the ^13^C-^19^F doublet were calculated as the matrix elements of the relaxation superoperator corresponding to the left and the right component of the ^13^C doublet. For the τ_c_-dependent series, BRW calculations were repeated in Spinach 2.10 under an isotropic rotational diffusion model on a grid of τ_c_ = 10, 15, 20, 25, 30, 40, 50, 70, 100 ns at a fixed 600 MHz (^1^H) spectrometer field. For each molecule (5-FU and 2Cl-4S), the CSA principal values/axes and the C-F internuclear vector for the active ^13^C–^19^F pair were taken from Gaussian16 shielding tensors and optimized geometries; the one-bond coupling was set to ^1^J_CF_ = 265 Hz for both systems to isolate CSA–DD effects (remote ^1^H were omitted in this sweep as they do not alter the qualitative trends). Relaxation superoperators were generated in the lab frame and projected onto single-quantum manifolds corresponding to TROSY, anti-TROSY, and a scalar-decoupled pathway (modeled by suppressing ^1^J_CF_ during relaxation propagation). Reported R_2_ values are the diagonal elements of the projected superoperators, and results are plotted separately for the ^19^F_C_ and ^13^C_F_ channels over 10–100 ns.

### Expression and purification of MBP

BL21(DE3) cells were transformed and plated overnight. The following morning, individual colonies were picked and used to inoculate 1-liter LB cultures (without glucose supplementation) for each construct. Cultures were grown at 37°C until they reached an OD_600_ of approximately 0.5, then induced with 1 mM IPTG. Following induction, cells were incubated for an additional 3 hours at 37°C and harvested by centrifugation at 8,000 × g for 10 minutes and stored at −80 °C until further use. The cell pellets were resuspended in 80 ml of lysis buffer containing 50 mM HEPES (pH 8), 150 mM NaCl, 1:1000 β-mercaptoethanol (BME), DNase, and half a protease inhibitor tablet. Cells were lysed by sonication while stirring on ice using a 5 s on/10 s off cycle at 35% amplitude for a total of 5 minutes, repeated twice. The lysate was clarified by centrifugation at 24,424 × g for 60 minutes, and the supernatant was filtered. The supernatant is pre-treated to ensure optimal binding to the amylose resin, then loaded onto a pre-equilibrated amylose resin column (Cytiva) using the Cytiva AKTA GO system. The flow rate is kept slow to maximize protein-resin interaction. The wash buffer consisted of 50 mM HEPES (pH 8), 150 mM NaCl, and 5 mM TCEP. The columns were then washed with 10 x column volumes of buffer. The protein is generally eluted using a buffer containing 10 mM maltose (50 mM HEPES (pH 8), 150 mM NaCl, and 5 mM TCEP). The elution profile is monitored by the AKTA system, and protein-containing fractions are collected based on absorbance at 280 nm.

### Expression and purification of NTSR1

The enNTS1 plasmid was transformed into *E. coli* BL21(DE3) cells and plated overnight on LB agar supplemented with 100 μg/mL carbenicillin at 37 °C. For expression, single colonies were used to inoculate overnight LB starter cultures containing 100 μg/mL carbenicillin, which were incubated at 37 °C and 220 RPM. These were used to seed 1 L cultures of M9 minimal media, supplemented with 100 μg/mL carbenicillin and 0.3% (w/v) glucose. The cultures were grown at 37 °C and 220 RPM until they reached an OD600 of approximately 0.15, after which the temperature was lowered to 16 °C. Once the OD600 reached approximately 0.6, protein expression was induced by the addition of 1 mM IPTG, and the cultures were incubated for ∼21 hours at 16 °C and 220 RPM. Finally, the cells were harvested by centrifugation at 4,000 × g and stored at −80 °C until further use.

Cell pellets derived from 2 liters of culture were thawed on ice and resuspended in 40 mL of buffer containing 100 mM HEPES, 400 mM NaCl, 20% glycerol at pH 8.0, supplemented with an EDTA-free protease inhibitor tablet (Roche), 100 mg lysozyme, and 10 mg DNase. The mixture was gently rocked at 4 °C for 30 minutes. After incubation, the suspension was sonicated on ice to ensure thorough cell lysis. After sonication, CHAPS (0.6% w/v), CHS (cholesterol hemisuccinate, 0.12% w/v), and DM (n-decyl-β-D-maltopyranoside, Anatrace, 1% w/v) solution were added. The resulting solubilization mixture was gently mixed at 4 °C for 2 hours to extract membrane proteins. Finally, the sample was clarified by centrifugation at 20,000 × g for 30 minutes at 4 °C. Solubilized enNTS1-containing supernatant was incubated with pre-equilibrated TALON resin (Cytiva) in buffer containing 25 mM HEPES, 10% glycerol, 300 mM NaCl, and 0.05% DDM (n-dodecyl-β-D-maltopyranoside, Anatrace) (pH 8.0) at 4°C for 15 minutes. The protein bound TALON resin was washed sequentially with two different buffers: the first wash buffer contained 25 mM HEPES, 10% glycerol, 500 mM NaCl, 0.05% DDM, 4 mM ATP, and 10 mM MgCl_2_ (pH 8.0), and the second wash buffer contained 25 mM HEPES, 10% glycerol, 350 mM NaCl, 0.05% DDM, and 10 mM imidazole (pH 8.0).

After washing, enNTS1 was eluted using a buffer containing 25 mM HEPES, 10% glycerol, 500 mM NaCl, 0.03% DDM, and 350 mM imidazole (pH 8.0). The elution profile is monitored by the AKTA system, and protein-containing fractions are collected based on absorbance at 280 nm. The eluate was incubated with 3 mg of 3C protease at 4°C for 16 hours to cleave off the MBP and muGFP tags. The cleaved receptor was concentrated using a 30 kDa MWCO centrifugal concentrator at 3,500 × g and then diluted 10-fold with SP ion-exchange equilibration buffer (20 mM HEPES, 10% glycerol, 0.03% DDM, pH 7.4).

This diluted sample was loaded onto a SP ion-exchange column (Cytiva) pre-equilibrated with the same buffer using a Cytiva AKTA GO system. The column was washed with SP wash buffer containing 20 mM HEPES, 10% glycerol, 250 mM NaCl, and 0.03% DDM (pH 7.4) until the absorbance at 280 nm stabilized. A 1 ml Ni²⁺-NTA column was connected in tandem with the SP column, and the receptor was eluted using SP elution buffer composed of 20 mM HEPES, 10% glycerol, 1 M NaCl, 0.03% DDM, and 25 mM imidazole (pH 7.4).

The resulting eluted protein was again concentrated using a 30 kDa MWCO concentrator at 4,000 × g and injected onto a GE S200 Increase size-exclusion chromatography (SEC) column equilibrated in NMR buffer (20 mM HEPES, 50 mM NaCl, 0.03% DDM, pH 7.5). Fractions containing the properly folded and purified enNTS1 were pooled, concentrated to 100–300 μM, and stored at −80°C until further use.

### Cysteine-Specific Labeling and kinetic analysis of MBP with Fluorinated Probe

Protein samples were prepared in a labeling buffer composed of 50 mM HEPES (pH 8.0), 150 mM NaCl, 5 mM EDTA, and 5 mM TCEP. BTFMA and 19F probes were each prepared at 0.5 M stock concentration in d-DMSO. For labeling, 25 μL of the respective probe solutions were added to each sample to achieve a final probe concentration of 25 mM, resulting in each probe-to-protein molar ratio and 5% final DMSO concentration. Reactions were carried out at room temperature in 1.5 mL Eppendorf tubes.

The addition of the probes caused a drop in pH to approximately 4–5. To restore neutral pH, 1–2 μL of 10 M NaOH was carefully added while mixing to prevent localized pH spikes. Final pH values of all reactions were confirmed to be between 7.2 and 7.4 using a ^19^F NMR-compatible micro pH electrode. Cloudiness was observed in the samples post-probe addition, likely due to low aqueous solubility of the fluorinated probes.

For kinetic analysis, aliquots were taken at 0, 15, 30, 45, 90-, 150-, 240- and 360-minutes post-reaction initiation. Each aliquot (1 μL) was quenched in 19 μL of 1% formic acid and immediately frozen at −20 °C for subsequent ESI-MS analysis. Following labeling, samples were clarified by centrifugation at 20,000 × g for 10 minutes and protein concentrations were determined from the supernatants. The labeled proteins were subjected to overnight dialysis at 4 °C using 10 kDa MWCO dialysis cassettes against 4 L of dialysis buffer (8000-fold volume excess). On the following day, samples were diluted five-fold and subsequently concentrated to a final volume of 500 μL. For NMR analysis, samples were prepared in buffer containing 50 mM HEPES (pH 7.0), 100 mM NaCl, 5% D2O, and 50 μM each of DSS and TFA as internal standards.

### Cysteine-Specific Labeling of NTSR1 with Fluorinated Probe

NTSR1(Q301C) was labeled with a cysteine-reactive ^19^F probe under conditions optimized for membrane protein stability. Purified receptor at 50 µM was prepared in NMR buffer containing 20 mM HEPES (pH 7.5), 50 mM NaCl, and 0.03% (w/v) DDM. Labeling reactions were carried out in a total volume of 300 µL by adding the probe from a concentrated stock solution to a final concentration of 5 mM. Samples were gently mixed and incubated overnight at 4 °C to allow complete modification of the engineered cysteine. After the reaction, unreacted probe and low–molecular-weight species were removed by size-exclusion chromatography (SEC) using the NMR buffer.

### NMR Spectroscopy

All NMR experiments were performed on MBP protein samples labeled with fluorinated probes. Unless otherwise specified, all spectra were acquired at 298 K in the designated NMR buffer. All 2D NMR datasets were processed using NMRPipe, a widely used UNIX-based spectral processing system^47^, and subsequently analyzed with Poky^48^. 1D NMR spectra were processed using MestReNova software (Mnova, Mestrelab Research)^49^.

### ^1^H NMR Experiments

1H NMR spectra were collected on a a Bruker Avance III HD 600 MHz spectrometer equipped with a 5 mm TCI CryoProbe. FID signals were acquired using a recycling delay of 2.7 seconds, and an acquisition time of 1.7 seconds. Each spectrum consisted of 128 scans, generating a free induction decay (FID) with 32768 complex points. All 1H NMR data were processed with a cosine-squared window function using MestReNova.

### 1D ^19^F NMR Experiments

1D ^19^F spectra were collected on a Bruker Avance III HD 600 MHz spectrometer equipped with a 5 mm triple resonance inverse CryoProbe (TCI, 1H tunable to 19F) located at the Integrated Molecular Structure Education and Research Center (IMSERC), Northwestern University. The 19F channel was set as the observation channel without proton or carbon decoupling to measure the native fluorine signal. The spectral width was set to 90 ppm with the carrier frequency centered at −145 ppm. 4096 complex points were acquired in the direct dimension, corresponding to an acquisition time of 0.045 seconds. A total of 4096 scans were collected with a 0.545 second recycling delay.

### 1D ^19^F with ^13^C Decoupling Measurements

To selectively suppress ^19^F -^13^C scalar couplings, additional 1D ^19^F NMR spectra were acquired with ^13^C decoupling applied during acquisition. The ^13^C carrier frequency was set to 150 ppm to effectively cover the aliphatic carbon region. This approach simplifies the ^19^F spectra by removing ^19^F -^13^C multiplet patterns, thereby improving the analysis of chemical shifts. Аll other acquisition parameters, including spectral width of 90 ppm, carrier frequency at –145 ppm, number of complex points of 4096, acquisition time of 0.045 seconds, number of scans of 4096, and recycle delay of 0.545 seconds, were identical to those used in the non-decoupled ^19^F experiment.

### 1D ^19^F with ^1^H, ^13^C Decoupling Measurements

For selective observation of ^19^F signals without scalar couplings to either ^1^H or ^13^C, one-dimensional ^19^F NMR spectra were acquired with simultaneous ^1^H and ^13^C decoupling. The ^13^C decoupling carrier was set at 151.5 ppm to ensure effective broadband decoupling of fluorine-labeled carbons. All other acquisition parameters, including a spectral width of 90 ppm, 4096 complex points, an acquisition time of 0.045 seconds, and 4096 scans, were identical to those used in the corresponding non-decoupled ^19^F NMR experiment.

### 1D ^13^C NMR Measurements

1D ^13^C NMR spectra were acquired using a standard gradient-enhanced pulse sequence (zggpwg) on a Bruker Avance III HD 600 MHz spectrometer. The spectral width was set to 80 ppm with the ^13^C carrier frequency centered at 140 ppm. 819 complex points and an acquisition time of 0.34 seconds. A relaxation delay of 1.34 seconds and 1024 scans were used.

### 1D ^19^F Chemical Shift Anisotropy (CSA) NMR Experiments

The ^19^F Chemical Shift Anisotropy with ^1^H decoupling solid-state NMR experiments were recorded on a 600 MHz Bruker Avance III HD NMR Spectrometer with a 1.6 mm HFXY Phoenix probe. The frequency of ^1^H was 599.77 MHz and the frequency of ^19^F is 564.43 MHz. The 90 degree pulse width of ^19^F nuclei was calibrated to 5.25 us. The magic angle spinning rate was set to 8000 Hz. 50 kHz SPINAL ^1^H heteronuclear decoupling was applied during acquisition. The recycle delay was set to 1,200 s per user direction. The MAS experiments averaged 24 transients. All spectra were referenced indirectly to the ^1^H resonance of adamantane (*δ*(1H) = 1.82 ppm) using the IUPAC convention. The MAS spectroscopy was deconvoluted by Dmfit^33^ and chemical shift anisotropy tensors was extracted.

### 2D ^19^F-^13^C and ^13^C- ^19^F out-and-stay TROSY Experiments

2D ^19^F-^13^C TROSY correlation spectra were recorded to characterize the fluorinated probe in poteins. All spectra were acquired on the same 600 MHz Bruker Avance III HD spectrometer with a 5 mm TCI CryoProbe tuned to ^19^F/^13^C frequencies. The ^1^H channel was tuned to ^19^F.

The spectral width in the indirect ^13^C dimension was set to 5 ppm, while the direct 19F dimension was set to 5 ppm. 1024 complex points were collected in the indirect ^13^C dimension (acquisition time = 686 ms), and 256 complex points were collected in the direct ^19^F dimension (acquisition time = 45 ms). The ^13^C and ^19^F carrier frequencies were centered at 150 ppm and −137.25 ppm, respectively. A total of 80 scans were collected per experiment with a 1.5 second recycling delay. Complementary 2D ^13^C –^19^F TROSY correlation spectra were acquired using the same 600 MHz Bruker Avance III HD spectrometer equipped with a 5 mm TCI CryoProbe configured for ^19^F /^13^C operation. These spectra were recorded with ^13^C as the direct detection channel and ^19^F as the indirect dimension. The spectral width in the indirect ^19^F dimension was set to 9.845 ppm, while the direct ^13^C dimension was set to 3.000 ppm. A total of 2048 complex points were collected in the indirect ^19^F dimension (acquisition time = 184 ms), and 128 complex points were acquired in the direct ^13^C dimension (acquisition time = 141 ms). The ^19^F and ^13^C carrier frequencies were centered at −137.0 ppm and 154.5 ppm, respectively. Each spectrum was acquired with 288 scans and a 2-second recycle delay.

### Data Processing

All 2D NMR data were processed using NMRPipe and analyzed with Poky. 1D spectra, including ^19^F and ^1^H data, were processed using Bruker TopSpin® (version 4.1.4) and MestReNova.

### ^19^F T_1_ Relaxation Measurements (^19^F_C_)

The longitudinal relaxation times (T_1_) of the fluorine nuclei (^19^F_C_) were measured using a 1D inversion recovery pulse sequence. Experiments were performed on Bruker Avance III HD 600 MHz spectrometer with a 5 mm TCI CryoProbe.

A experiment was acquired with the following variable inversion recovery delays: 10, 100, 200, 600, 1200, and 2000 ms.

Each experiment was recorded with 128 scans and a recycled delay of 5.16 s to ensure full relaxation. The 19F carrier frequency was set to −145 ppm to selectively excite the fluorine resonance.

Data were processed using MestReNova and OriginPro software and fitted to a mono-exponential recovery curve to extract the T_1_ values.

The longitudinal relaxation times (T_1_) were determined by fitting the signal intensities obtained from the inversion recovery experiments to the following mono-exponential recovery equation:

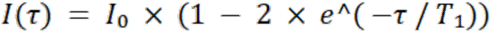

where I(τ) is the observed signal intensity at a given inversion recovery delay τ, I_0_ is the equilibrium signal intensity, and T_1_ is the longitudinal relaxation time to be determined.

### ^19^F T_2_ Relaxation Measurements (^19^F_C_)

The transverse relaxation times (T_2_) of ^19^F nuclei (^19^F_C_) were determined using a CPMG (Carr–Purcell–Meiboom–Gill) sequence to minimize magnetic field inhomogeneity effects.

Measurements were performed on the same Bruker Avance III HD 600 MHz spectrometer. The ^19^F carrier frequency was set to −145 ppm.

A series of experiments were recorded with variable CPMG echo times, and the decay of signal intensities was fitted to a mono-exponential decay function to determine T_2_ values.

The transverse relaxation times (T_2_) were determined by fitting the decay of signal intensities to a mono-exponential function according to the equation:

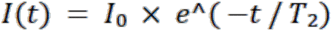

where I(t) is the signal intensity at time t, I_0_ is the initial signal intensity, and T_2_ is the transverse relaxation time.

### ^13^C T_1_ Relaxation Measurements (^13^C_F_)

The longitudinal relaxation times (T_1_) of ^13^C nuclei directly bonded to fluorine (^13^C_F_) were measured using a 1D inversion recovery pulse sequence. Experiments were carried out on a Bruker Avance III HD 600 MHz spectrometer, equipped with a cryogenically cooled probe to enhance sensitivity. A experiment was performed with the following inversion recovery delays: 50, 150, 300, 600, 1000, 1400, and 2200 ms. Each experiment was acquired with 256 scans and a recycle delay of 5.22 s. The ^13^C carrier frequency was set at 155 ppm to selectively excite the ^13^C_F_ resonance.

### ^13^C T_2_ Relaxation Measurements (^13^C_F_)

The transverse relaxation times (T_2_) of ^13^C nuclei (^13^C_F_) were measured using a ^13^C CPMG sequence. The experiments were conducted on the same Bruker Avance III HD 600 MHz spectrometer with a cryoprobe. The ^13^C carrier frequency was set at 155 ppm to observe the ^13^C_F_ signals. A series of variable CPMG echo times were applied, and the decay of the signal intensity was monitored.

### Mass Spectrometry (Intact MS)

Intact mass spectrometry analysis was performed at the Integrated Molecular Structure Education and Research Center (IMSERC), Northwestern University, utilizing the Agilent 6220A Time-of-Flight LC–MS system. This high-resolution instrument (resolving power ∼10,000 at m/z 1522) is coupled with liquid chromatography (LC) for online sample cleanup, followed by electrospray ionization (ESI) to introduce the analyte into the mass spectrometer. Protein samples were analyzed under native-like LC conditions, and mass spectra were acquired capturing the charge-state envelope. Deconvolution to neutral molecular masses was performed using a Maximum Entropy algorithm.

### Samples for viscosity/τ_c_ estimation, and NMR measurement

MBP K34C/R354C was expressed and purified by standard procedures and stoichiometrically labeled with the cysteine-reactive 2Cl-4Cl probe. Final NMR samples contained 20% (w/w) glycerol, 50 mM HEPES buffer, 150 mM NaCl, and 5% D2O at pH 7.5. Spectra were acquired at 25 °C, 15 °C, and 5 °C with ≥10 min equilibration at each temperature. Dynamic viscosities for 20% (w/w) glycerol–water were obtained from the Cheng correlation^24^ (temperature-dependent values for w = 0.20) and used directly in subsequent analyses; the resulting viscosities were ≈1.51/2.00/2.77 mPa·s at 25/15/5 °C, respectively. Rotational correlation times (τ_c_) were then estimated using a standard Stokes–Einstein–Debye scaling referenced to MBP in water at 25 °C (τ_c_ ≈ 25 ns; water viscosity 0.89 mPa·s), yielding τ_c_ ≈ 42/58/84 ns for 20% glycerol at 25/15/5 °C. Two complementary experiments were used: ^19^F →^13^C TROSY (^19^F -observed with ^13^C editing) and ^13^C →^19^F TROSY (^13^C -observed with ^19^F editing). Carriers were centered on the 2Cl-4Cl ^19^F and the directly bonded ^13^C. Acquisition parameters were matched across temperatures. Data were processed identically; FWHM values were obtained from Lorentzian fits to peak cross-sections and S/N as peak height over rms noise. Split components are annotated as P1/P1b/P1c (K34C) and P2/P2b/P2c (R354C).

All spectra were acquired on the same 600 MHz Bruker Avance III HD spectrometer with a 5 mm TCI CryoProbe tuned to ^19^F/^13^C frequencies. The ^1^H channel was tuned to ^19^F.

The spectral width in the indirect ^13^C dimension was set to 5 ppm, while the direct ^19^F dimension was set to 2 ppm. 2048 complex points were collected in the indirect ^13^C dimension (acquisition time = 1.372 s), and 128 complex points were collected in the direct ^19^F dimension (acquisition time = 56 ms). The ^13^C and ^19^F carrier frequencies were centered at 153 ppm and −137.65 ppm, respectively. A total of 288 scans were collected per experiment with a 0.5 second recycling delay.

And the spectral width in the indirect ^19^F dimension was set to 10 ppm, while the direct ^13^C dimension was set to 3 ppm. 2048 complex points were collected in the indirect ^19^F dimension (acquisition time = 0.18 s), and 128 complex points were collected in the direct ^13^C dimension (acquisition time = 0.14 s). The ^13^C and ^19^F carrier frequencies were centered at 153.5 ppm and −137 ppm, respectively. A total of 288 scans were collected per experiment with a 2 second recycling delay.

Both 2D experiments were acquired with 50% nonuniform sampling (NUS) in the indirect dimension; spectra were reconstructed by compressed sensing with identical reconstruction parameters across temperatures.

### DFT-based calculation and visualization of ^13^C and ^19^F shielding tensors

Molecular structures of fluorinated aromatic compounds were obtained from density functional theory (DFT) geometry optimizations. Optimized Cartesian coordinates were exported in Protein Data Bank (PDB) format using GaussView and used as structural input for tensor visualization. Nuclear magnetic shielding tensors for the selected ^13^C and ^19^F nuclei were extracted from the corresponding Gaussian output files as full 3 × 3 tensors expressed in the molecular Cartesian frame. The tensors were symmetrized by averaging off-diagonal elements to yield physically meaningful second-rank shielding tensors.

Visualization of molecular structures and chemical shielding anisotropy tensors was performed using TensorView for MATLAB^50^. The PDB structures were loaded into TensorView, and the symmetrized shielding tensors were assigned to the corresponding nuclear coordinates. Shielding tensors were displayed exclusively as ellipsoids, representing the principal axis system obtained by tensor diagonalization, with TensorView applying the appropriate rotations internally. Visualization parameters, including tensor scaling and transparency, were kept consistent across all structures, and dipolar coupling tensors or vectors were omitted to focus on nucleus-specific shielding anisotropy. Final three-dimensional renderings were exported for figure preparation.

### Synthesis of ^13^C-2Cl-4Cl and 2Cl-4F (natural abundance)

#### General procedures

^13^C-bromoacetic acid was purchased from Cambridge Isotope Laboratories (Isotope.com). Selectfluor^TM^ (1-Chloromethyl-4-fluoro-1,4-diazoniabicyclo[2.2.2]octane bis(tetrafluoroborate)) and 2,4-dichloro-5-fluoropyrimidine (2Cl-4Cl, natural abundance) were purchased from Fluorochem Ltd (Fluorochem.co.uk). Pd/BaSO_4_ (5%) was purchased from ChemImpex, USA (Chemimpex.com). All other reagents and solvents were purchased from Fluorochem and Valerus Ltd (Valerus-bg.com) and used without additional purification, unless noted otherwise. NMR spectra were obtained on a Bruker AVIII 500 MHz instrument. Electrospray mass spectrometry was performed on Waters Micromass ZQ 2000. Flash chromatography was performed on a Waters Delta 600 system equipped with Teledyne Isco 340CF evaporative light scattering detector, using silica gel and C18 cartridges purchased from BGB Analytik (Bgb-analytik.com).

#### 2-Cyanoacetic-2-13C acid (1)

5.0 g ^13^C-bromoacetic acid (36 mmol) was dissolved in 16 ml water. A solution of 3 g Na_2_CO_3_ (28 mmol) in 11 ml water was added in portions until the pH reached 9. 2.37 g KCN (36 mmol) was dissolved in 8 ml water and added drop-wise. The mixture was stirred for 4 hours at 80°C, then at room temperature for 24 hours, and acidified to pH 1 with 4 ml concentrated HCl. The solvent was removed under reduced pressure. The residual water was removed by dissolving the product in t-BuOH and removing the solvent under reduced pressure. The resulting solid was sonicated in 50 ml diethyl ether for 4 hours, filtered off and washed with diethyl ether. The combined Et_2_O extracts were evaporated at 60°C to yield 3.15 g of crude **1** (36 mmol, quantitative), which was used in the next step without further purification. ^1^H NMR (CDCl_3_/CD_3_OD, 500 MHz) δ 3.51 (d, *J* =136.4 Hz, 2H, *H*_2_C^13^C) ppm. ^13^C NMR (CDCl_3_/CD_3_OD, 125 MHz) δ 24.6 (*C*H_2_), 113.4 (d, *J* = 62.9, *C*N), 165.6 (d, *J* = 59.0, *C*OOH) ppm. LRMS (ESI): *m/z* calcd. for C_2_^13^CH_3_NO_2_ [M-H]^−^ : −85.54; found: −85.20.

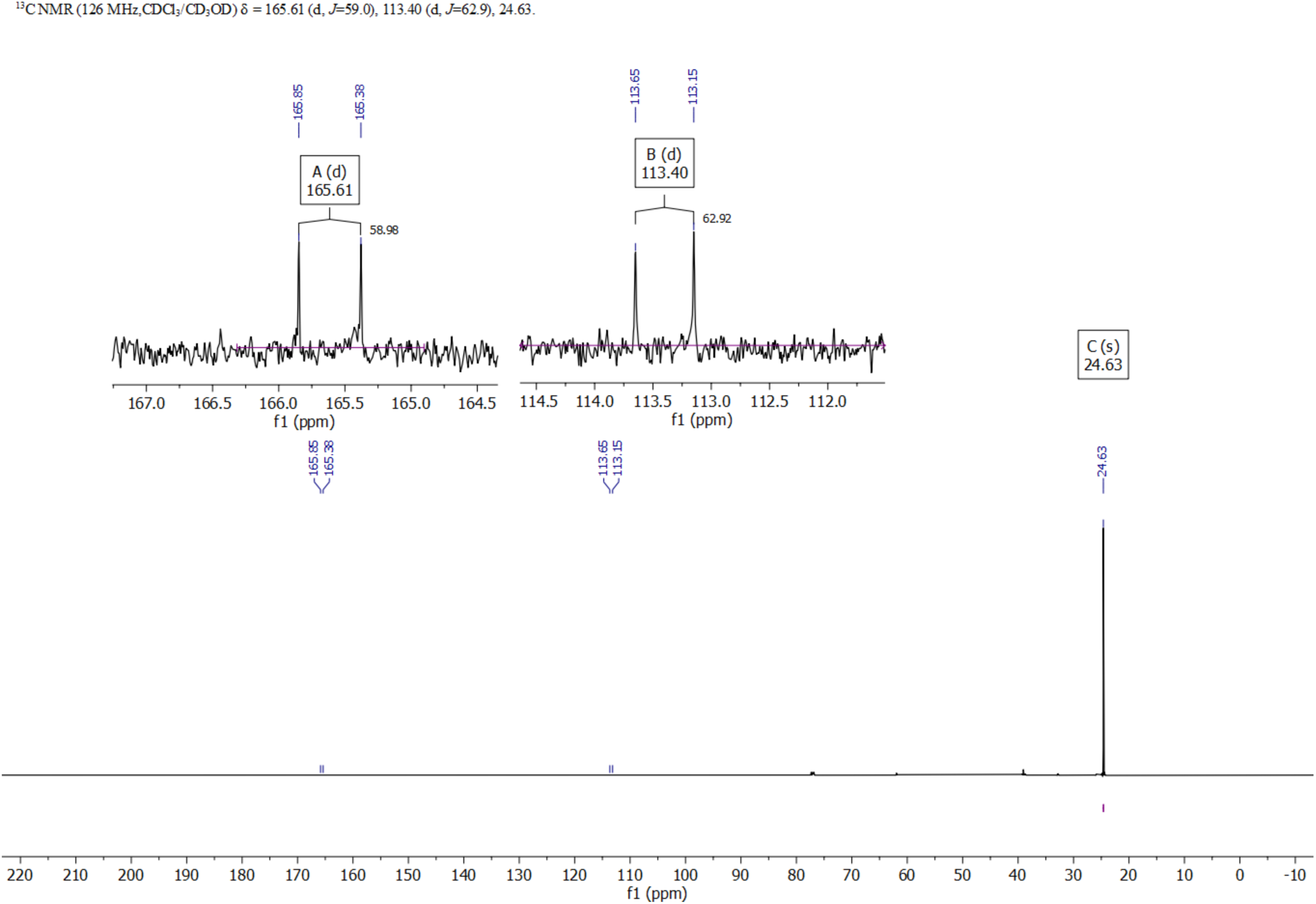

#### N-Carbamoyl-2-cyanoacetamide-2-^13^C (2)

1.57 g (18.2 mmol) of cyanoacetic acid **1** was dissolved with 1.31 g (21.8 mmol) urea in 11 ml acetic anhydride. The mixture was stirred at 90°C. After 10 minutes, a thick white precipitate formed. After 2 hours, 12 ml of water was added and the mixture was allowed to cool while stirring, upon which additional precipitate formed. The solid was filtered off, washed with 4ml water and dried for 48 hours to give 2.33 g of cyanourea **2** (18.2 mmol, quantitative) as white solid. ^1^H NMR (DMSO-d_6_, 500 MHz) δ 3.90 (d, *J* = 136.8 Hz, 2H, *H_2_*C^13^C), 7.35 (s, 2H, N*H_2_*), 10.38 (s, 1H, N*H*) ppm. ^13^C NMR (DMSO-d_6_, 125 MHz) δ 26.79 (*C*H_2_), 115.2 (d, J = 62.6, *C*N), 153.0 (H_2_N*C*ONH), 165.0 (d, J = 51.3, HN*C*OCH_2_) ppm. LRMS (ESI): *m/z* calcd. for C_3_^13^CH_5_N_3_O_2_ [M-H]^−^ : −127.04; found: −127.25.

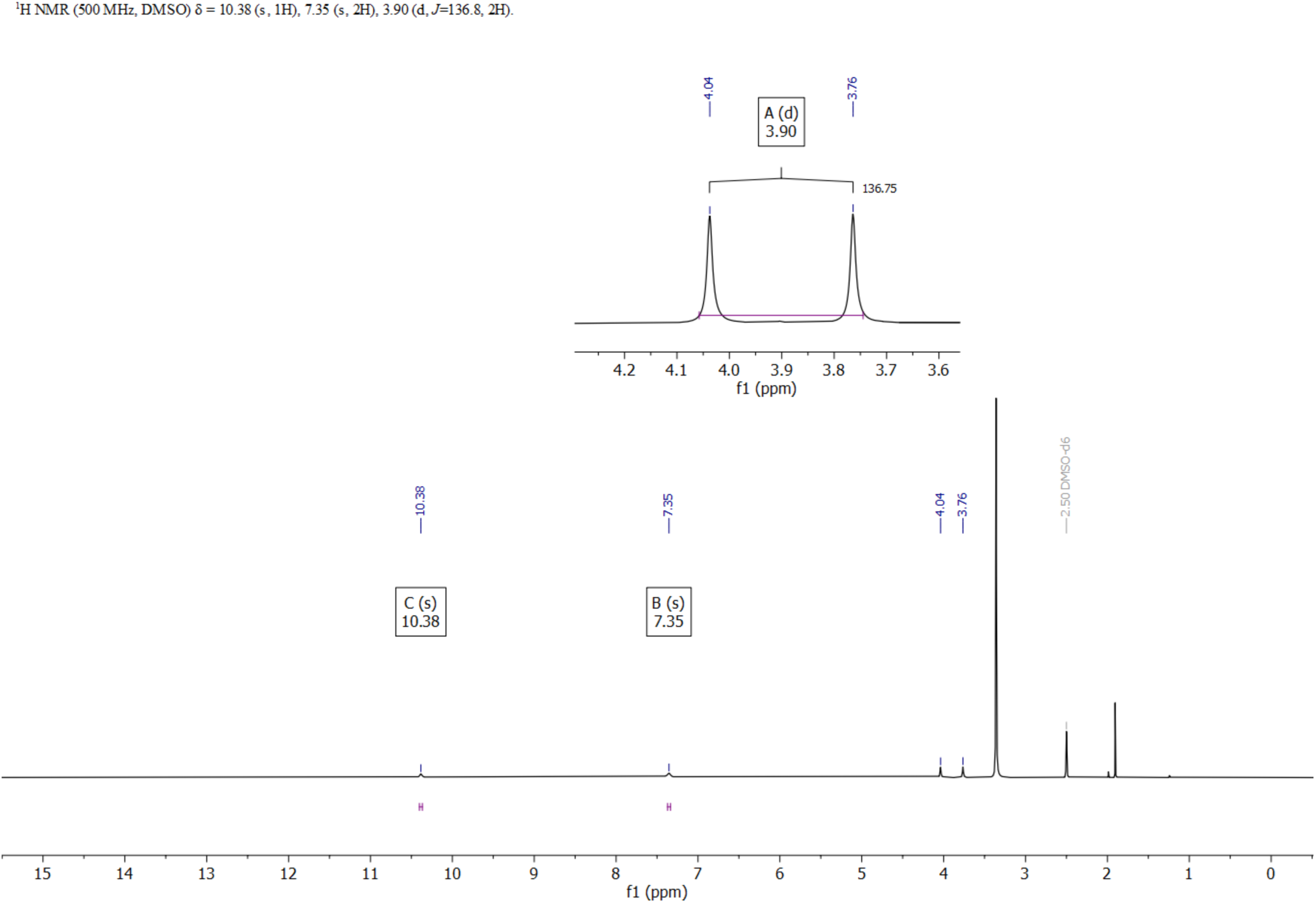

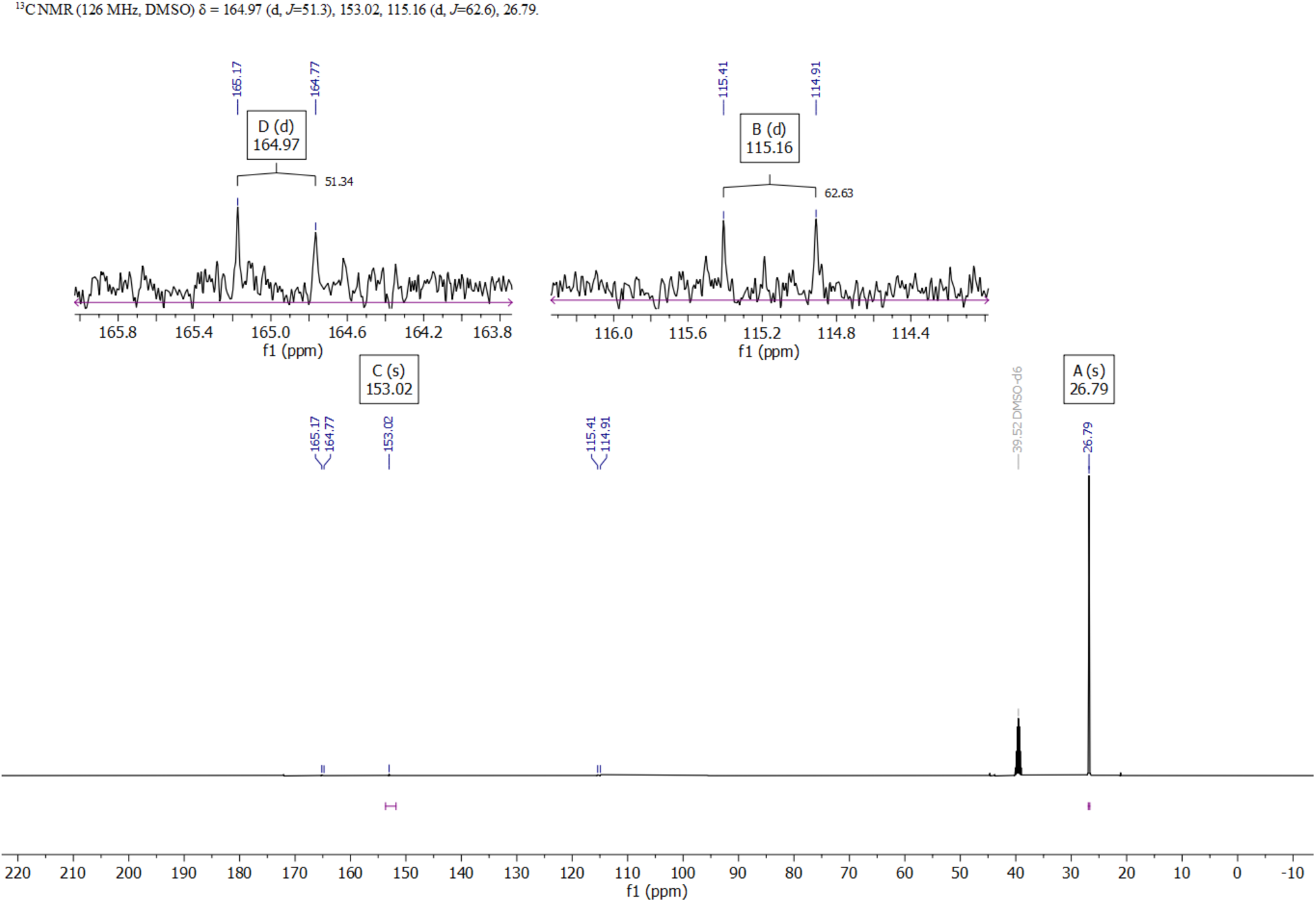

#### Pyrimidine-2,4(1*H*,3*H*)-dione-5-^13^C (5-^13^C-uracil) (3)

1.65 g Pd/BaSO4 (5%) was suspended in 50 ml of 50% acetic acid and the catalyst was activated by seven cycles of vacuum/hydrogen purge from a balloon. 2.33 g (18.2 mmol) of **2** was dissolved in 100 ml of boiling 60% aqueous acetic acid and added to the activated palladium catalyst. The mixture was stirred under a hydrogen atmosphere (balloon) at room temperature. After 60 hours the mixture was heated to 70°C and the catalyst was filtered off. The solvent was partially removed under reduced pressure until the formation of a white precipitate and placed at 4°C overnight. The precipitate was filtered off and dried on air overnight to give 1.38 g greyish solid. The filtrate was concentrated further to yield additional 0.16 g of product. The NMR spectra revealed residual acetic acid which was removed by suspending the product in 20 ml water and 10 ml toluene and stripping the solvents under reduced pressure to yield 1.40 g (12.4 mmol, 69%) of **3**. ^1^H NMR ( DMSO-d_6_, 500 MHz) δ 5.44 (dd, *J* = 174.85, 7.65 Hz, 1H), 7.38 (d, *J* = 4.78 Hz). ^13^C NMR (DMSO-d_6_, 125 MHz) δ 100.23, 142.24 (d, *J* = 65.9 Hz), 151.56, 164.38 (d, *J* = 64.8 Hz) ppm. LRMS (ESI): *m/z* calcd. for C_3_^13^CH_4_N_2_O_2_ [M-H]^−^ : −112.03; found: −112.21.

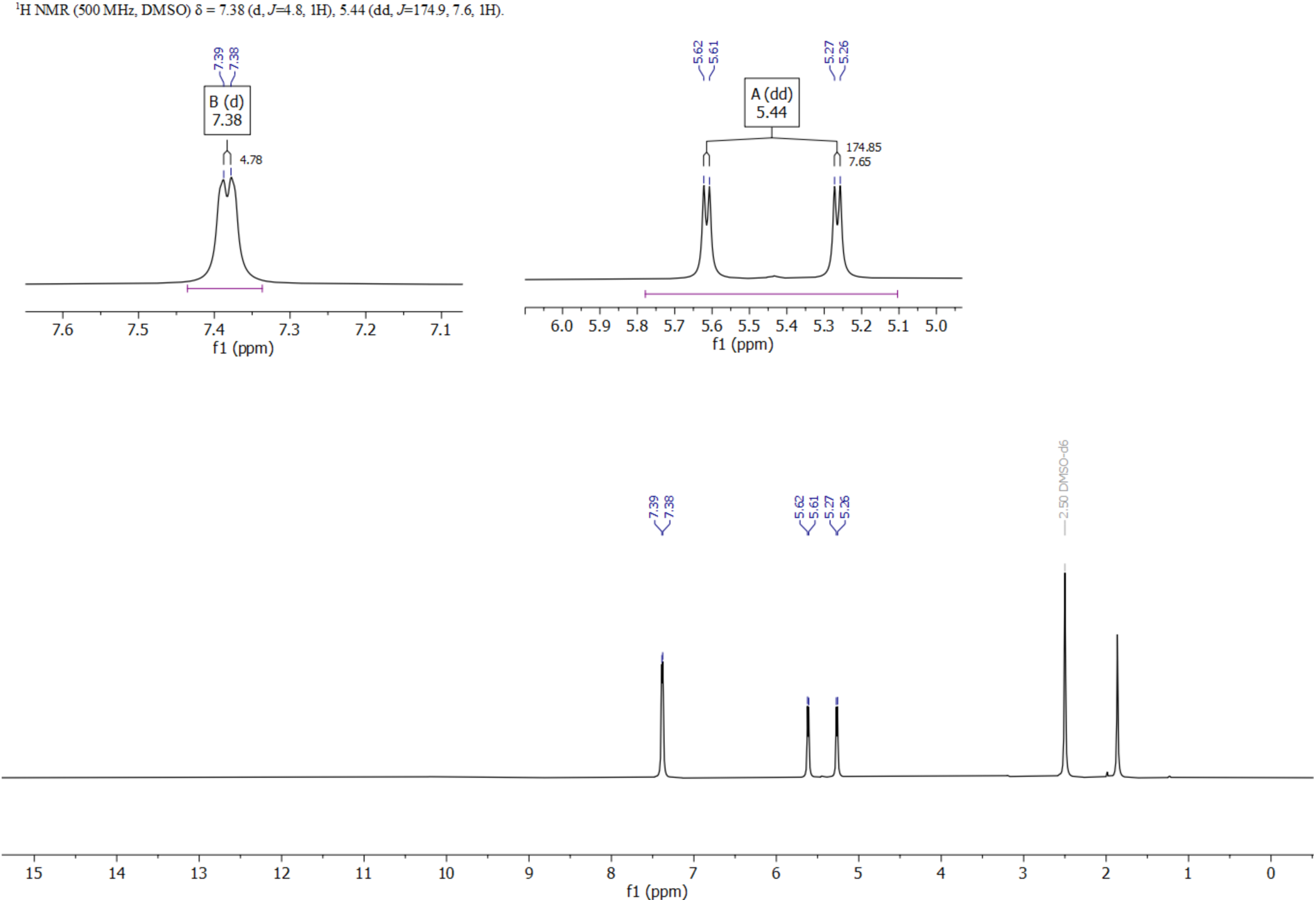

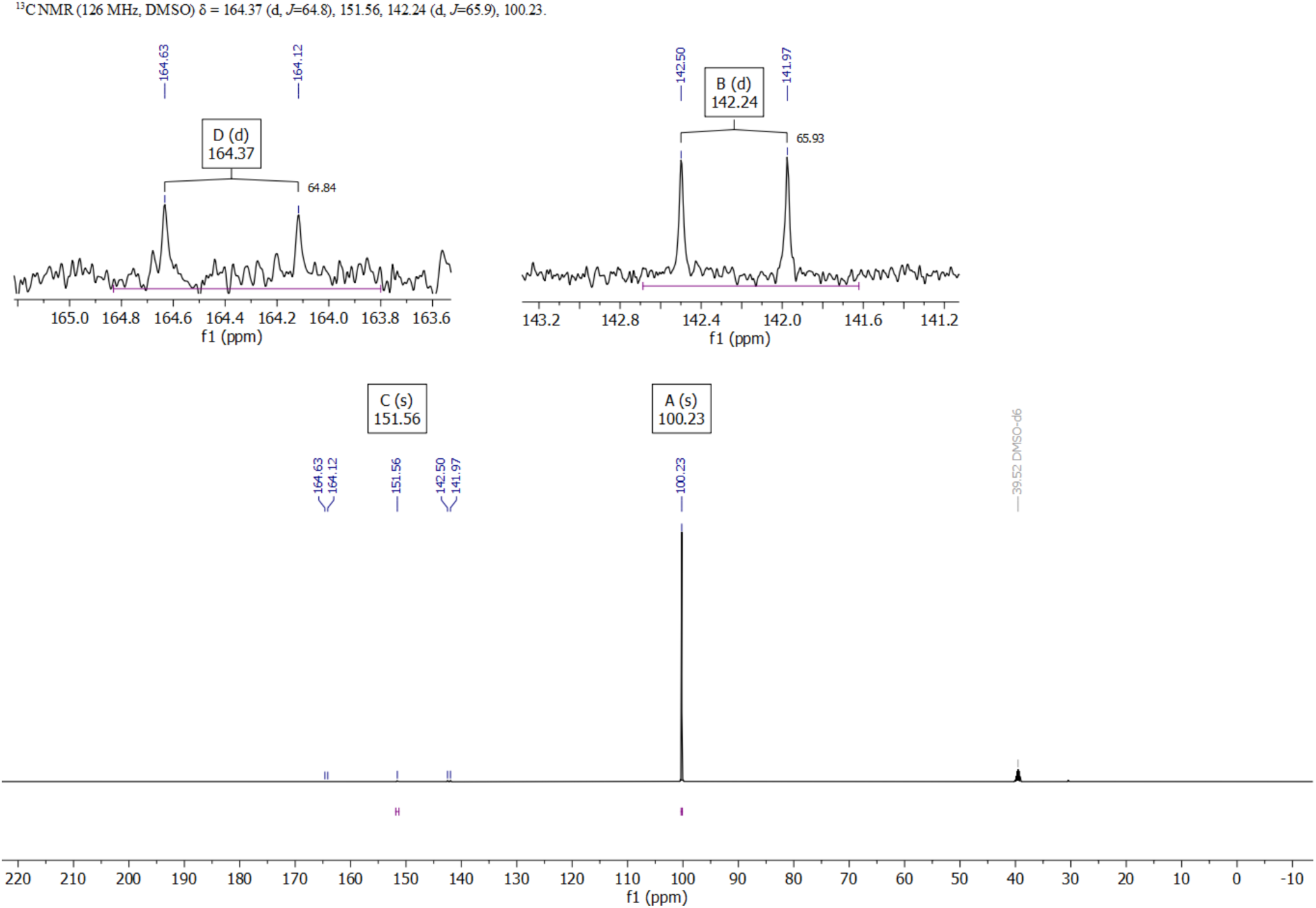

#### 5-Fluoro-6-hydroxydihydropyrimidine-2,4(1*H*,3*H*)-dione-5-^13^C (4)

4.78 g Selectfluor^TM^ (13.5 mmol) was added to a suspension of 1.40 g (12.4 mmol) of uracil **3** in 40 ml water. The mixture was stirred under argon atmosphere at 90°C for 4 hours. The heat was removed and a solution of 10.4 g NaBPh_4_ (30 mmol) in 140 ml water was added. The mixture was stirred for 2 hours and then placed at 4°C overnight. The precipitated solid was filtered and washed with water. The filtrate and water washes were combined and evaporated to dryness under reduced pressure, resulting in 4.50 g solid containing crude **4**, and a considerable amount of tetraphenylborate salts. This was used in the next step without purification.

#### 2,4-Dichloro-5-fluoropyrimidine-5-^13^C (6) (^13^C-2Cl-4Cl)

1.58 g of the crude **4** was dissolved in 15.6 g POCl_3_ (100 mmol). 1.70 g dimethylaniline (14.0 mmol) was added dropwise and the mixture was heated under reflux for 2 hours, then allowed to cool to room temperature overnight. The reaction mixture was poured onto 100 g ice, 50 ml water and 100 ml diethyl ether and stirred until the ice had melted. The organic phase was separated, and the aqueous phase was extracted three times with 100 ml of diethyl ether. The combined ether extracts were dried over anhydrous Na_2_SO_4_ and the solvent was carefully stripped under reduced pressure without a water bath. (*Note: The product sublimes easily and can be lost under reduced pressure, if heat is applied!)* The resulting oil was purified by flash chromatography with a gradient of 0 to 20% Et_2_O in hexanes to yield 225 mg **6** (1.34 mmol, 31 % from **3**, 21 % overall from 13C-bromoacetic acid). ^1^H NMR ( DMSO-d_6_, 500 MHz) δ 9.02 (dd, *<u>J</u>*_CH_= 1Hz, *<u>J</u>*_FH_ = 1Hz, 1H) ppm. ^13^C NMR (DMSO-d_6_, 126 MHz) δ 153.59 (d, *J* = 264.2 Hz). ^19^F NMR (DMSO-d_6_, 471 MHz) δ −135.25 (d, *J_CF_* = 264.4 Hz).

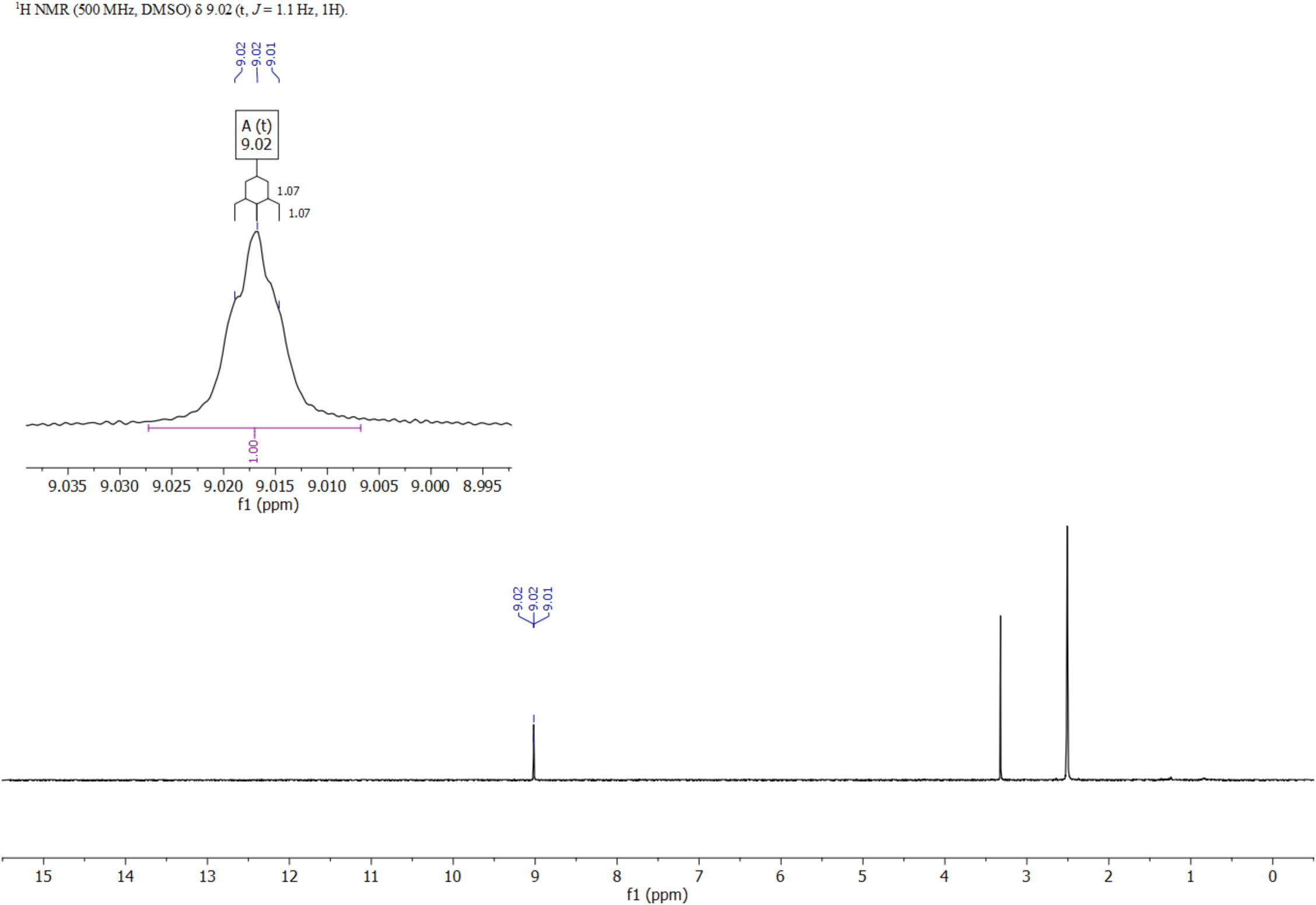

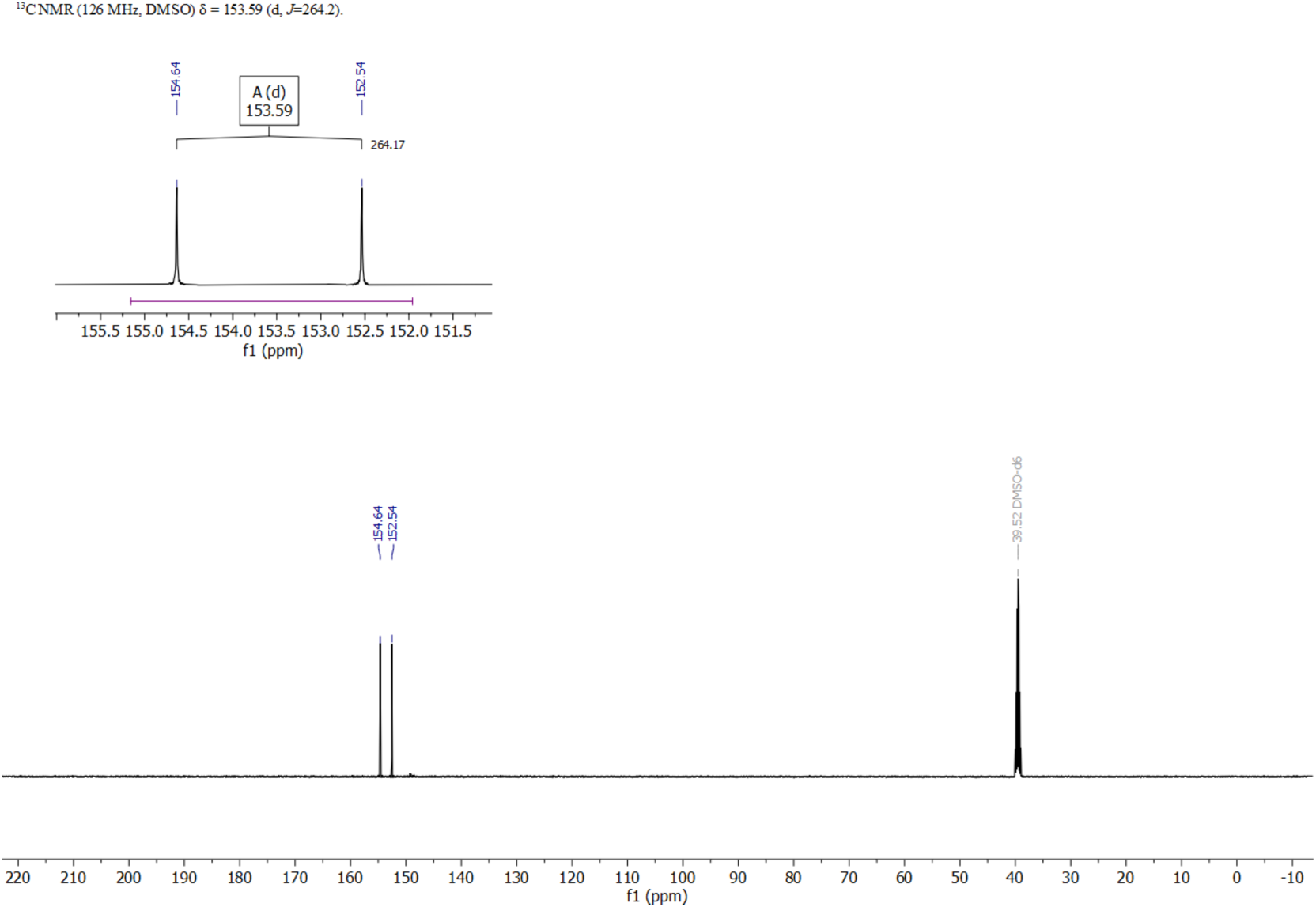

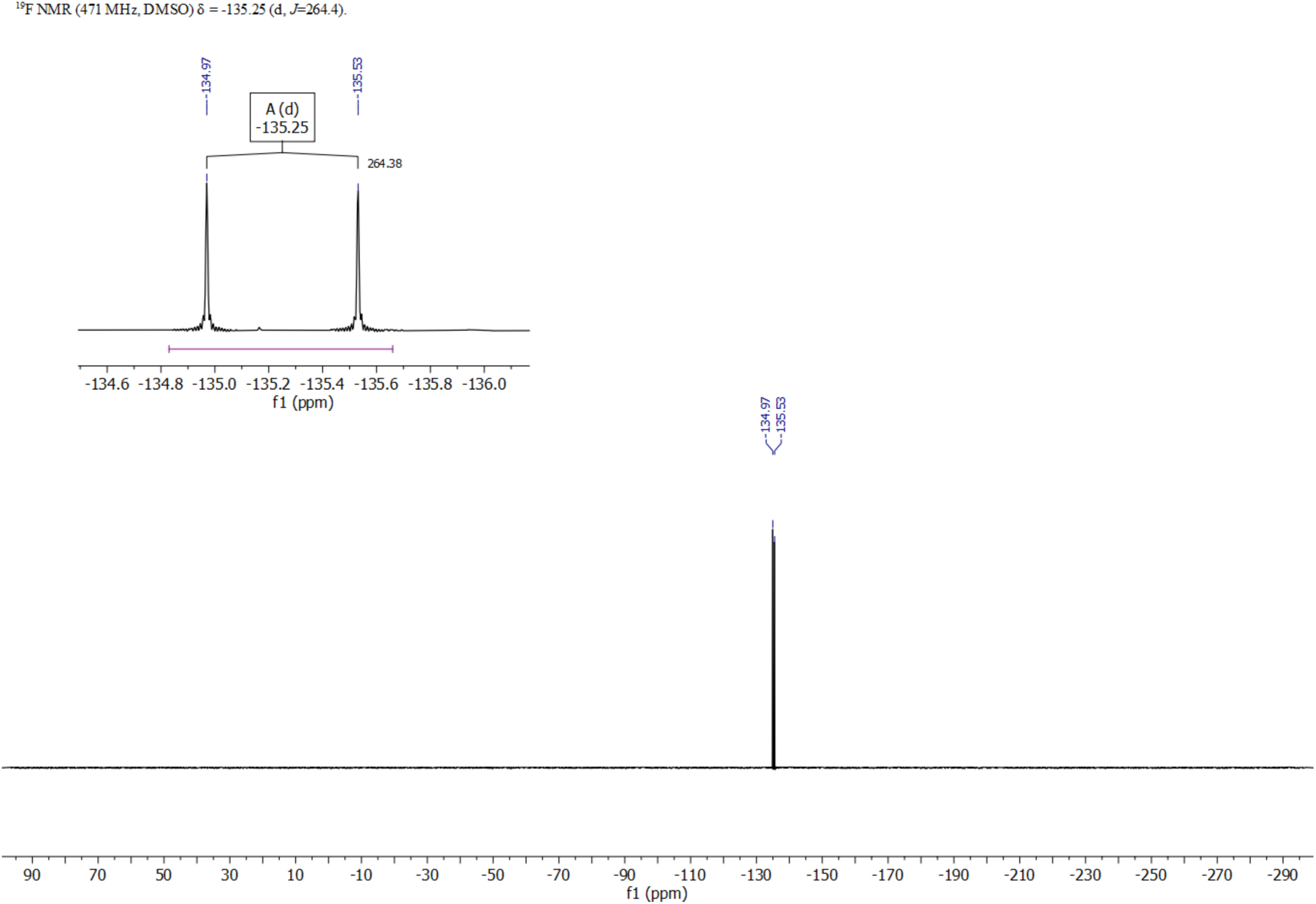

#### 2-Chloro-4,5-difluoropyrimidine (2Cl-4F)

0.38g ( 2.28 mmol) 2,4-Dichloro-5-fluoropyrimidine in 2ml DMSO (stored over 4A molecular sieves) was added to 0.52g (3.34 mmol) CsF (dried by heating for 5 minutes at 30 mbar and 200°C). The reaction mixture was stirred at 100°C in a screwcap vial. In 15 minutes a white precipitate formed. After 4 hours the heat was removed and the mixture was allowed to cool to room temperature overnight with continued stirring. The solution was loaded on a 12g silicagel cartridge and the product was eluted with a gradient of 0 to 25% Et_2_O in hexanes. The solvent was carefully stripped on a rotary evaporator without a water bath. (*Note: The product is extremely volatile and will be lost under reduced pressure, if any heat is applied!*)

^1^H NMR ( DMSO-d_6_, 500 MHz) δ 9.08 (d, *J*_FH_= 11.7Hz, *J*_CH_ = 2.0Hz, 1H) ppm. ^13^C NMR (DMSO-d_6_, 126 MHz) δ = 157.43 (dd, *J*=257.2, 14.4), 150.80 (dd, *J*=20.3, 4.0), 148.96 (d, *J*=23.6), 144.42 (dd, *J*=263.7, 20.2).. ^19^F NMR (DMSO-d_6_, 471 MHz) δ −76.57 (d, J = 23.6), −154.72 (d, J = 23.6)

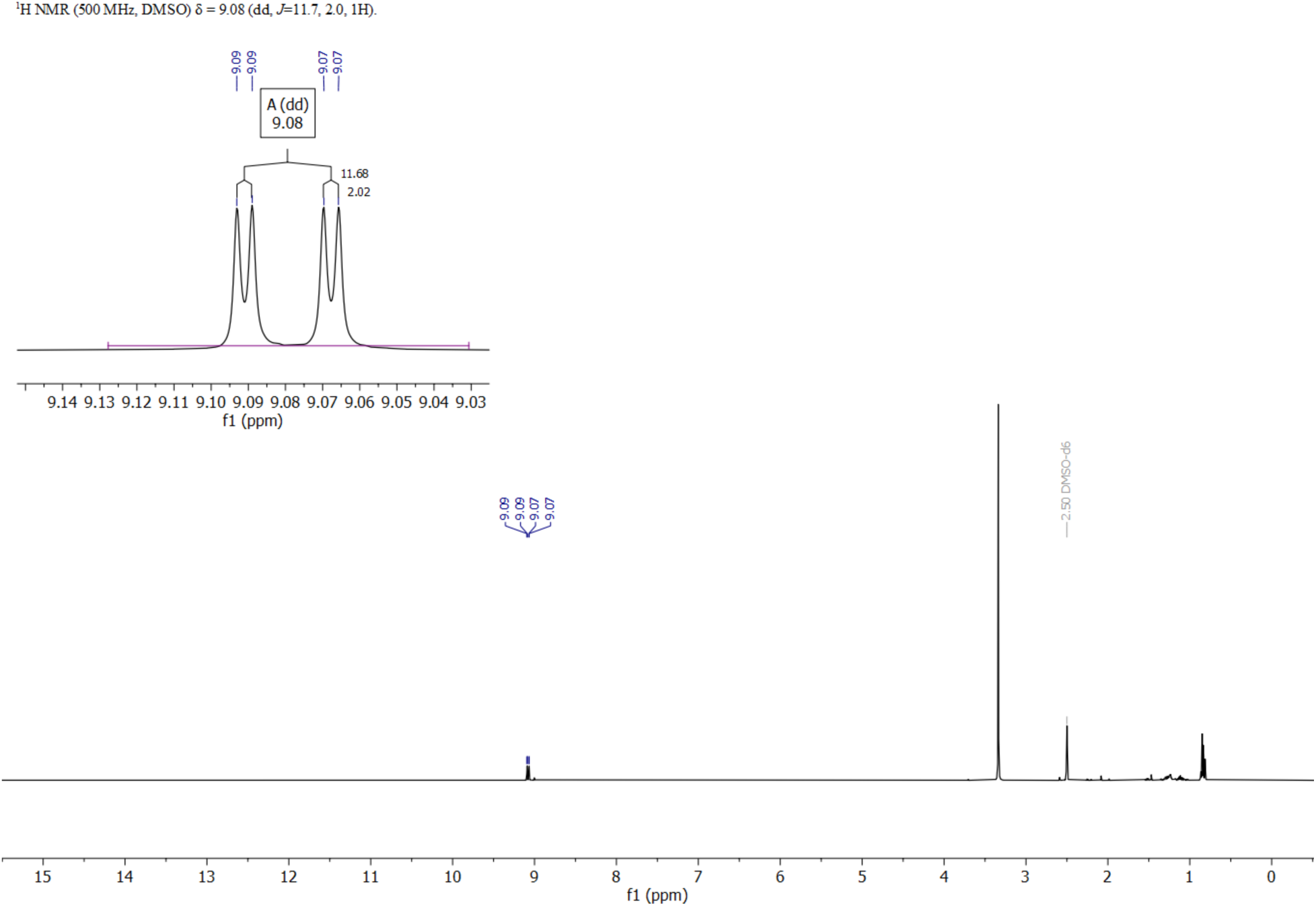

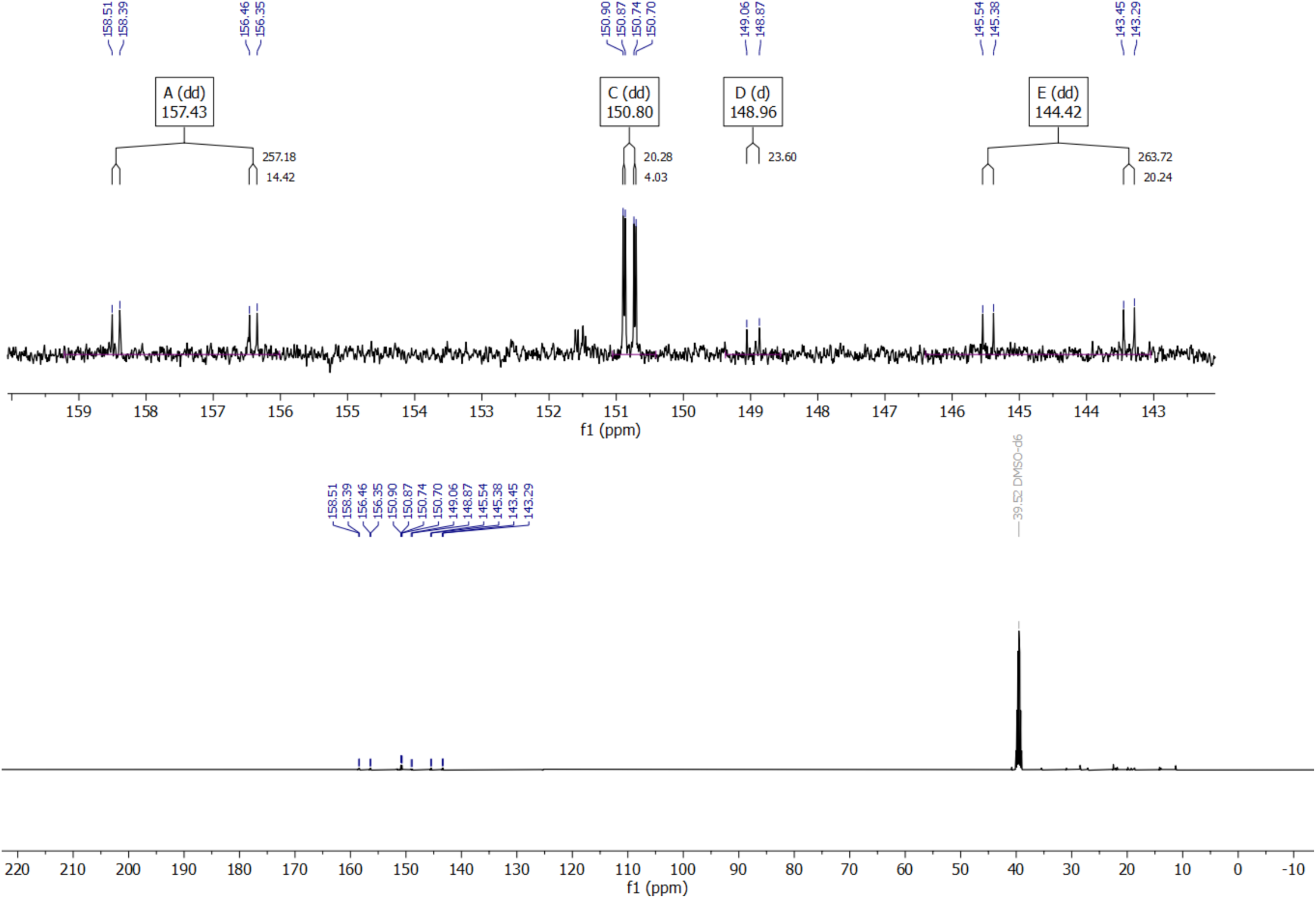

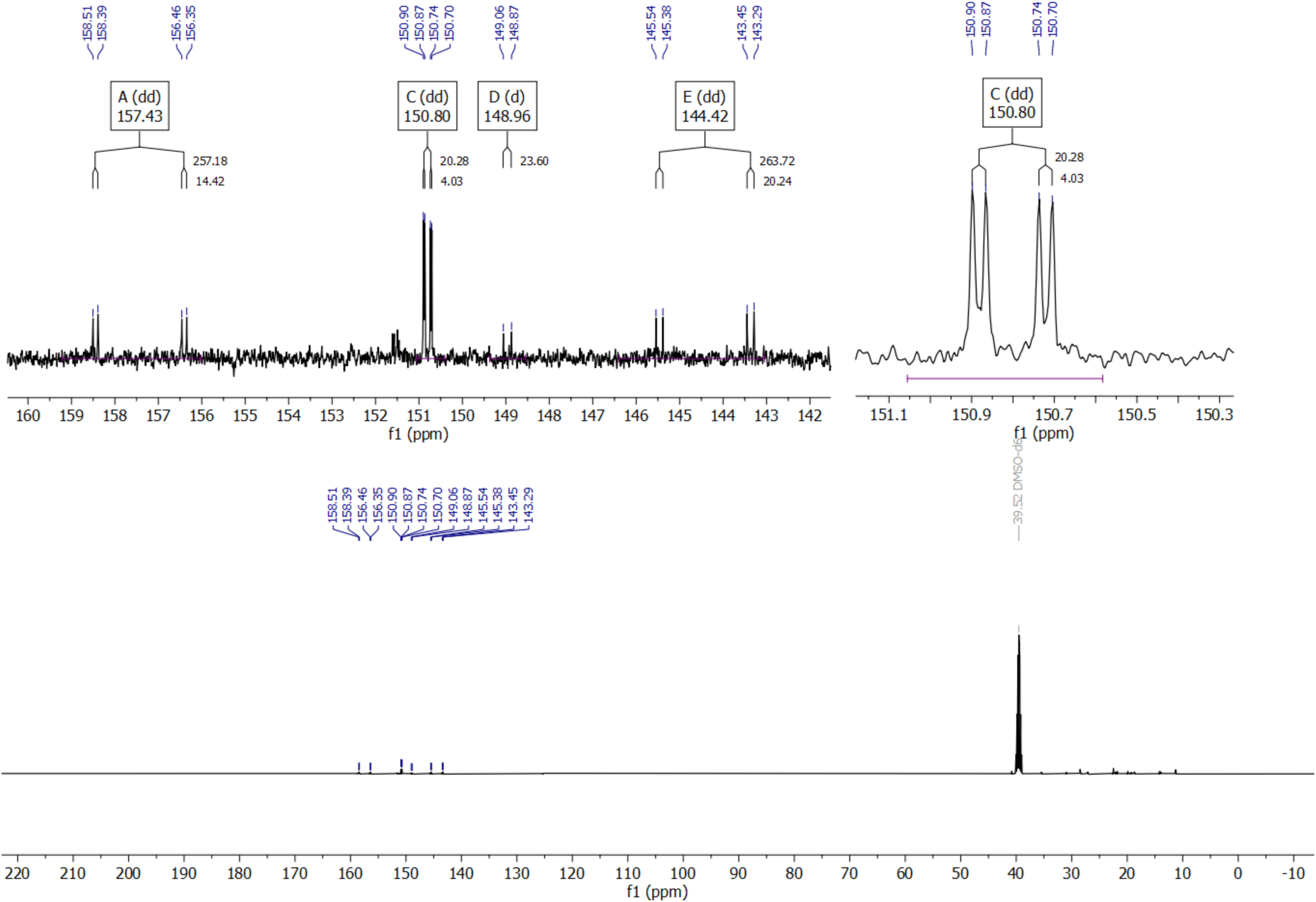

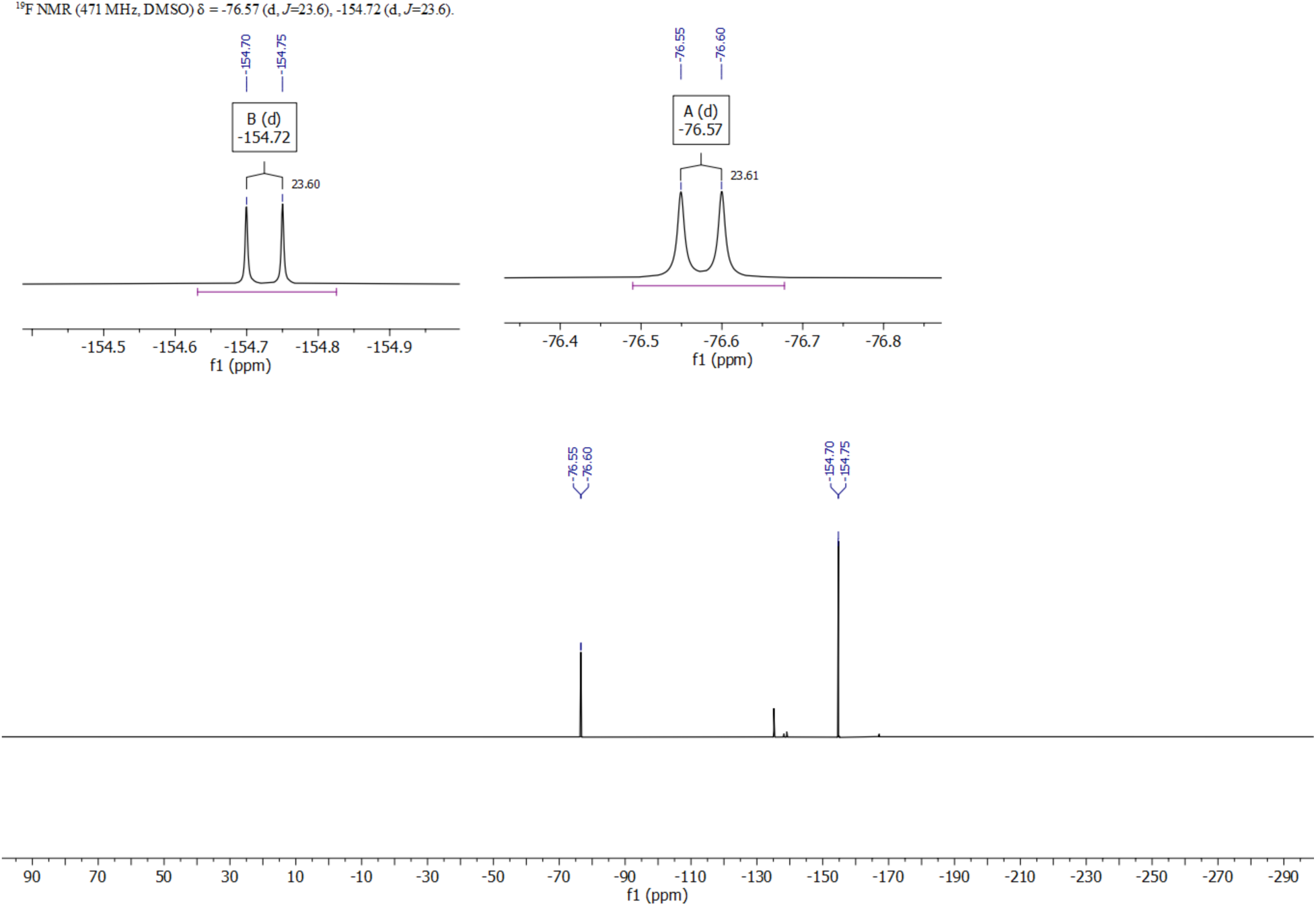

