## Supplementary Information for "Chemical control of CSA geometry enables relaxation-optimized ^19^F-^13^C NMR probes"

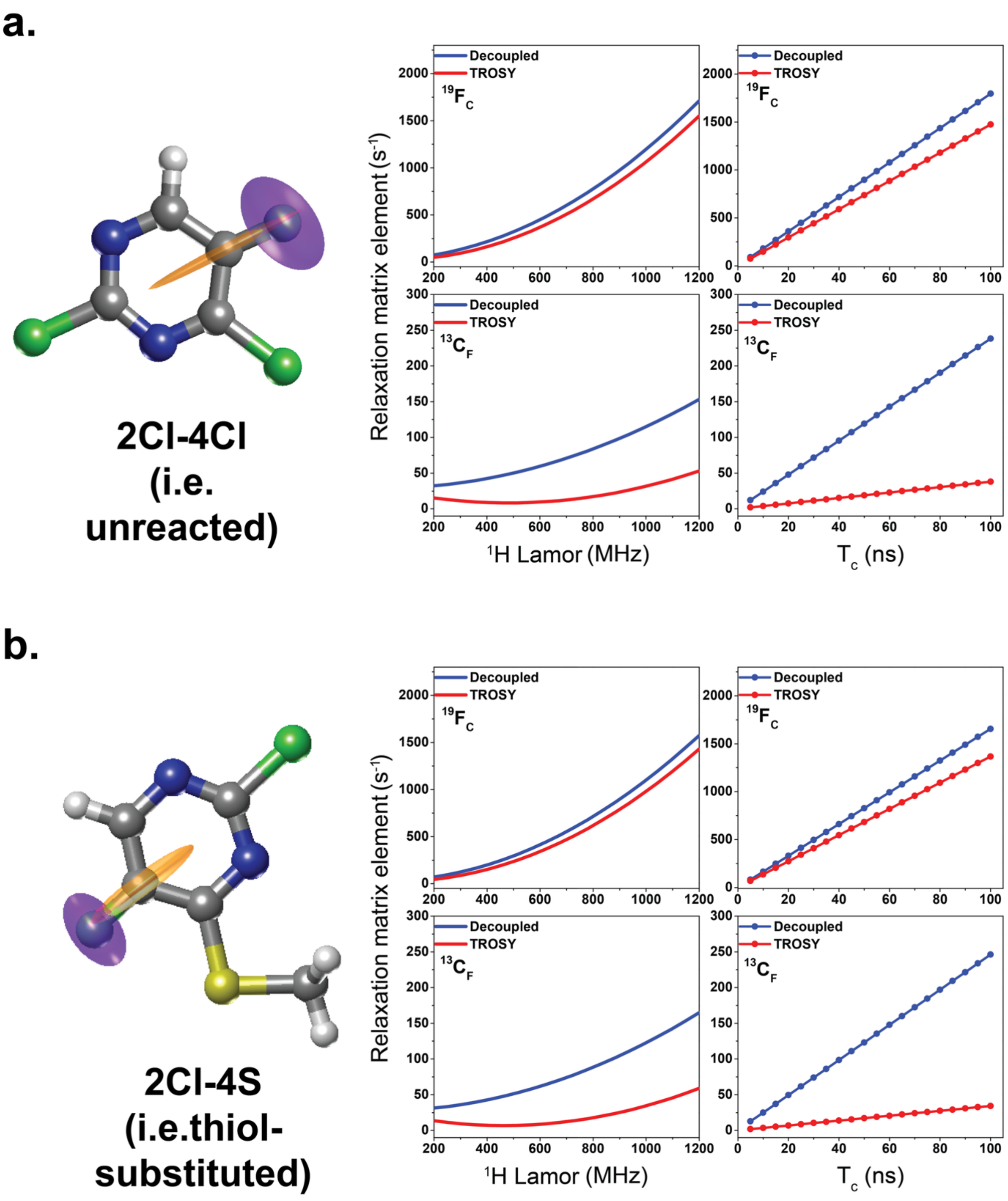


**Figure S1. Thiol-substitution is predicted to have little-to-no effect on CSA geometry and $^{19}$F-$^{13}$C TROSY performance.** Principal axes of the traceless $^{19}$F (purple) and $^{13}$C (orange) chemical shift anisotropy (CSA) tensors, calculated by density functional theory (DFT), overlaid on the molecular structures of 2Cl-4Cl (**a**) and 2Cl-4S (**b**). Corresponding Bloch-Redfield-Wangsness (BRW) simulations of TROSY (red) and decoupled (blue) transverse relaxation rates ($R_2$) for $^{19}F_C$/$^{13}C_F$ single-quantum coherences are shown as a function of $^1$H Larmor frequency (MHz) and rotational correlation time ($\tau_c$). All simulations were performed in the rigid limit ($S^2$ = 1) assuming isotropic tumbling with $\tau_c$ = 25 ns, providing an upper-bound, geometry-focused comparison across scaffolds rather than a quantitative reproduction of experimental relaxation rates.

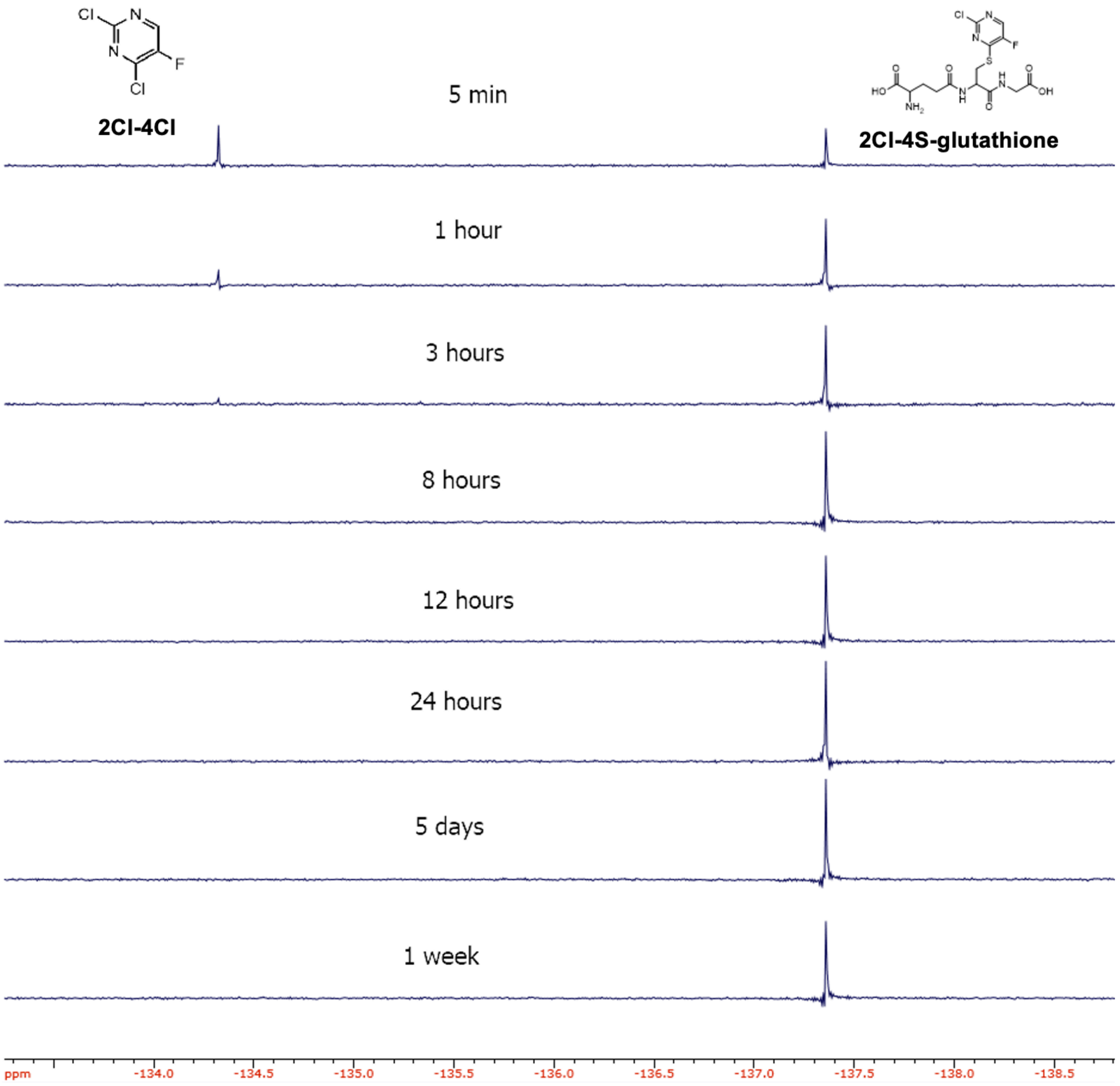


**Figure S2. Short-term stability of the 2Cl-4S-glutathione adduct at room temperature in 50 mM sodium phosphate buffer, pH 7.2.** $^{19}$F-spectra of 2 mM 2Cl-4Cl mixed with 2 mM glutathione at room temperature after 5 min to one week. Reacted is complete with 8 h with the 2Cl-4S-glutathione adduct exhibiting little-to-know degradation over a week.

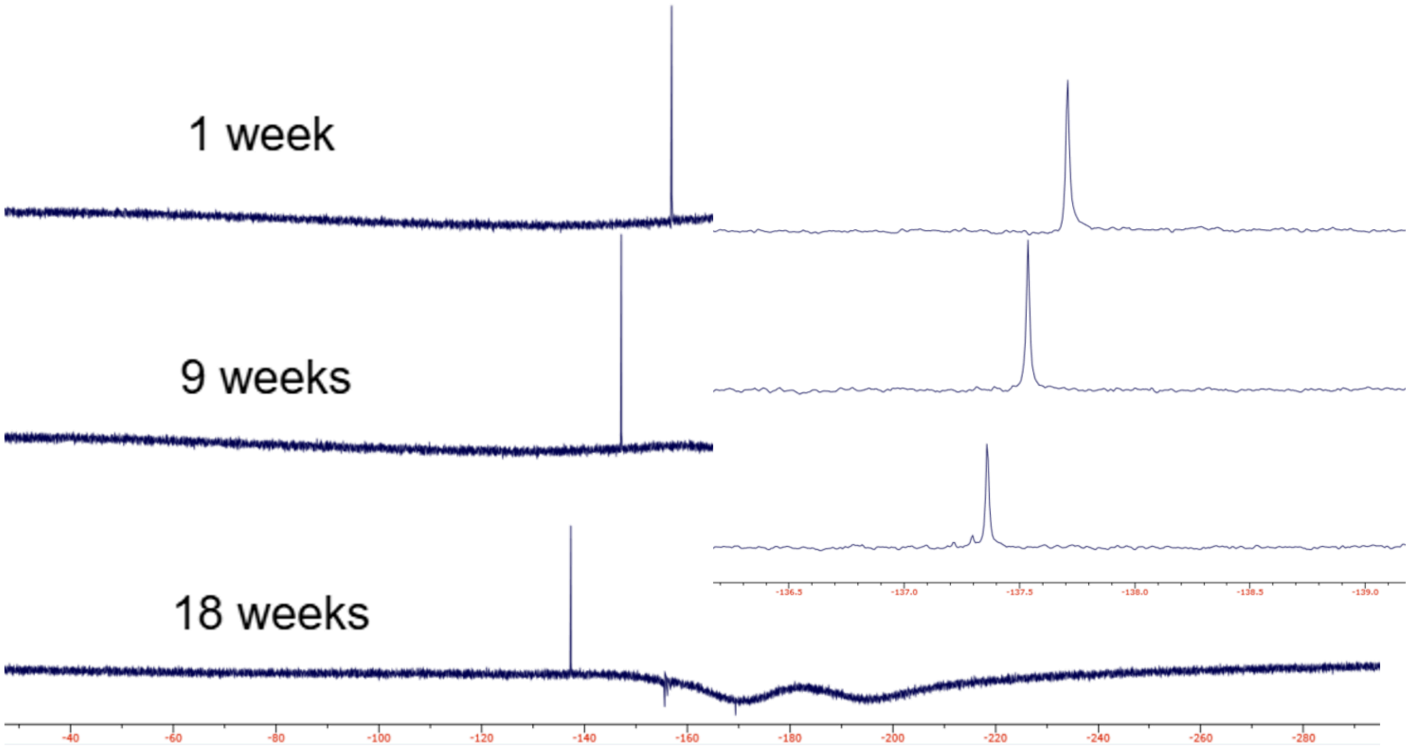


**Figure S3. Long-term stability of the glutathione-2Cl-4S adduct at room temperature in 50 mM sodium phosphate buffer, pH 7.2.** $^{19}$F-spectra of 2 mM 2Cl-4Cl mixed with 2 mM glutathione at room temperature after 1, 9 and 18 weeks.

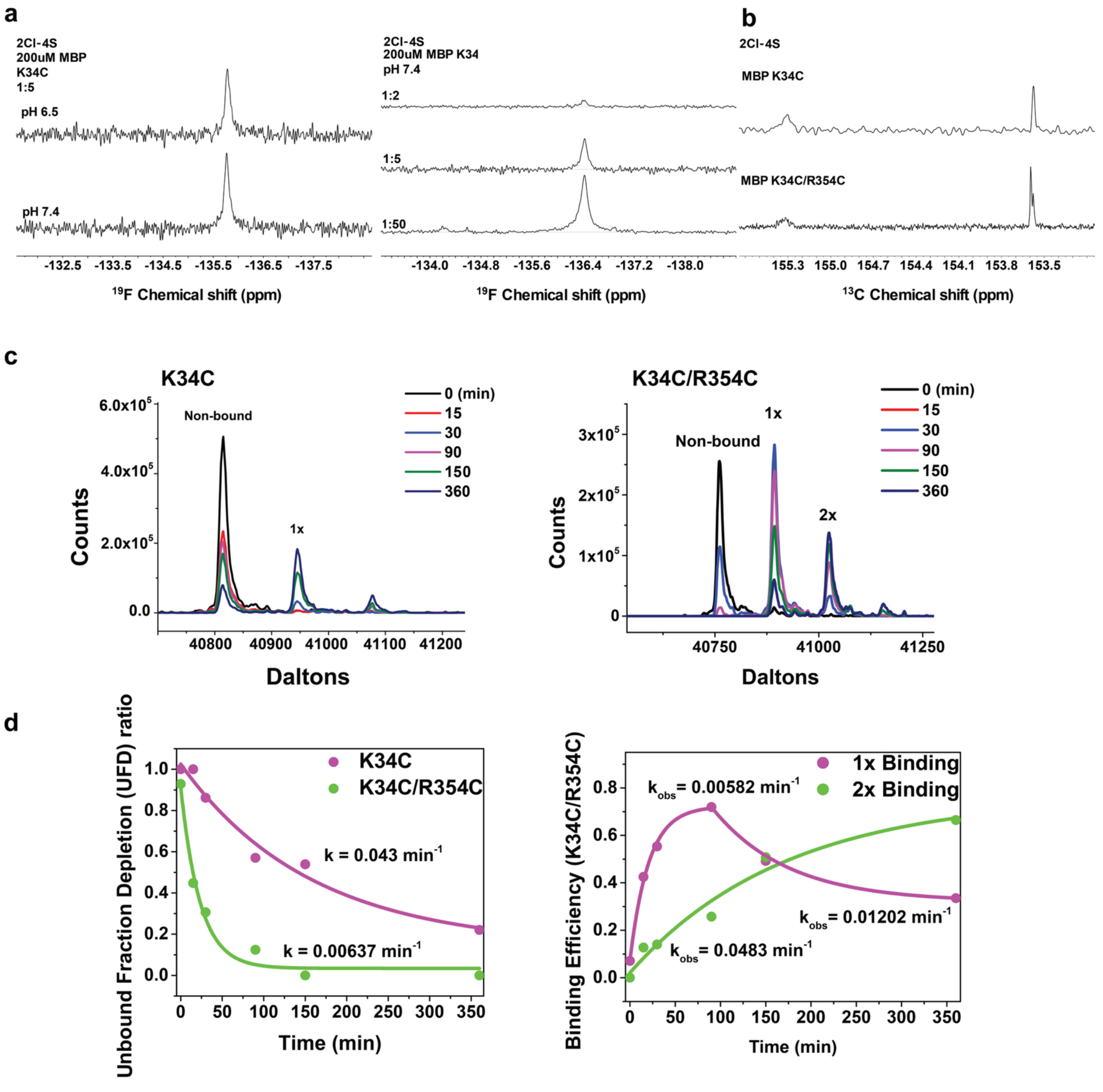


**Figure S4. NMR and Intact protein mass spectrometry analysis of 2Cl-4Cl conjugation to MBP variants. (a.)** $^{19}$F NMR spectra of 2Cl-4S-MBP(K34C) acquired under varying reaction conditions. **(a, *left*)** Effect of pH on conjugation efficiency, comparing spectra obtained at pH 6.5 and pH 7.4 for reactions performed at a protein:probe molar ratio of 1:5 (200 µM MBP). **(a, *right*)** Effect of protein-to-probe ratio at pH 7.4, showing $^{19}$F spectral profiles for 1:2, 1:5, and 1:50 molar ratios. The peak intensity and linewidth reflect the extent of probe conjugation and labeling homogeneity under each condition. **(b, *top*)** $^{13}$C spectra of MBP K34C showing two well-resolved resonances at 155.3 ppm (anti-TROSY) and 153.57 ppm (TROSY). **(b, *bottom*)** $^{13}$C spectrum of MBP(K34C/R354C) displaying an additional resonance at 153.60 ppm assigned to the R354C site overlapped with the K34C TROSY peak. **(c, *left*)** Time-resolved, deconvoluted mass spectra of MBP(K34C) reacted with 2Cl-4Cl, showing progressive appearance of the singly bound (1×) species from the non-bound protein. **(c, *right*)** Time-resolved, deconvoluted mass spectra of MBP(K34C/R354C) reacted with 2Cl-4Cl, displaying non-bound, singly bound (1×), and doubly bound (2×) species. **(d, *left*)** Time-dependent depletion of unmodified MBP (unbound fraction depletion, UFD) following reaction with 2Cl-4Cl. Labeling at K34C proceeds with a pseudo–first-order rate constant of $k = 0.043\ min^{-1}$, whereas the K34C/R354C double mutant shows slower depletion kinetics ($k = 0.0064\ min^{-1}$), reflecting sequential binding steps. **(d, *right*).** Reaction efficiency profiles of 2Cl-4Cl with MBP(K34C/R354C). Fitted rate constants ($k_{obs}$) indicating faster initial modification for the first site ($k_{obs} = 0.049\ min^{-1}$) followed by slower incorporation at the second site ($k_{obs} = 0.012\ min^{-1}$). These results highlight the regioselectivity and high efficiency of the thiol-specific chemistry of the 4-2Cl probe.

| Molecule | PAS | Eigenvalue (ppm) | Angle (deg) | Skew (κ) | Span (Ω, ppm) | $\sigma_{eff}$ (ppm) | $\lvert\sigma_{eff}\rvert/\Omega$ |
|---|---|---|---|---|---|---|---|
| **3-$^{19}F_{13C}$ Tyr ($^{19}F$)** | 1 | -61.72 | 83.17 | -0.47 | 146.30 | -23.40 | 0.16 |
| | 2 | -22.85 | 6.83 | | | | |
| | 3 | 84.57 | 89.97 | | | | |
| **3-$^{19}F_{13C}$ Tyr ($^{13}C$)** | 1 | -63.97 | 6.31 | 0.09 | 124.15 | -63.15 | 0.51 |
| | 2 | 3.80 | 83.69 | | | | |
| | 3 | 60.17 | 89.99 | | | | |
| **4-$^{19}F_{13C}$ Phe ($^{19}F$)** | 1 | -77.17 | 89.39 | 0.04 | 152.58 | 1.76 | 0.01 |
| | 2 | 1.76 | 0.81 | | | | |
| | 3 | 75.41 | 89.46 | | | | |
| **4-$^{19}F_{13C}$ Phe ($^{13}C$)** | 1 | -90.72 | 1.13 | 0.30 | 164.88 | -90.66 | 0.55 |
| | 2 | 16.55 | 89.54 | | | | |
| | 3 | 74.16 | 88.97 | | | | |
| **5-FU ($^{19}F$)** | 1 | -43.19 | 48.60 | -0.64 | 109.78 | -32.06 | 0.29 |
| | 2 | -23.41 | 41.40 | | | | |
| | 3 | 66.60 | 90.00 | | | | |
| **5-FU ($^{13}C$)** | 1 | -60.77 | 24.02 | 0.19 | 114.38 | -49.52 | 0.43 |
| | 2 | 7.17 | 65.98 | | | | |
| | 3 | 53.61 | 90.00 | | | | |
| **2Cl-4Cl ($^{19}F$)** | 1 | -48.61 | 82.36 | -0.69 | 126.15 | -29.28 | 0.23 |
| | 2 | -28.94 | 7.64 | | | | |
| | 3 | 77.54 | 90.00 | | | | |
| **2Cl-4Cl ($^{13}C$)** | 1 | -81.72 | 6.58 | 0.28 | 149.67 | -80.47 | 0.54 |
| | 2 | 13.77 | 83.42 | | | | |
| | 3 | 67.95 | 90.00 | | | | |

**Table S1. Principal values and orientations of $^{19}F$ and $^{13}C$ CSA tensors from DFT calculations.**
